# Parent-specific transgenerational immune priming enhances offspring defense – unless heat-stress negates it all

**DOI:** 10.1101/2024.03.29.587316

**Authors:** Ralf F Schneider, Arseny Dubin, Silke-Mareike Marten, Olivia Roth

## Abstract

Trans-generational immune priming (TGIP) adjusts offspring immune responses based on parental immunological experiences - a form of trans-generational plasticity predicted to be adaptive when parent-offspring environmental conditions match. In contrast, mismatches between environmental conditions negate those advantages, rendering TGIP costly when mismatched immunological offspring phenotypes are induced. Particularly maternal TGIP was thought to shape offspring immunological preparedness: mothers’ eggs contain more substance than sperm and, in viviparous species, pregnancy provides additional avenues for immune priming of developing offspring. The syngnathids’ (pipefishes and seahorses) unique male pregnancy provides an unusual perspective to the ecological relevance of TGIP in a system where egg production and pregnancy occur in different sexes. We simulated parental bacteria exposure in broad nosed pipefish, *Syngnathus typhle*, through vaccinations with heat-killed *Vibrio aestuarianus* before mating the fish to each other or control individuals. Resulting offspring were raised, and some exposed to *V. aestuarianus*, in a control or heat-stress environment, after which transcriptome and microbiome compositions were investigated. Transcriptomic TGIP effects were only observed in *Vibrio*-exposed offspring at control temperatures, arguing for low costs of TGIP in non-matching environments. Transcriptomic phenotypes elicited by maternal and paternal TGIP had only limited overlap and were not additive. Both transcriptomic responses were significantly associated to immune functions, and specifically the paternal response to the innate immune branch. TGIP of both parents reduced the relative abundance of the experimental *Vibrio* in exposed offspring, showcasing its ecological effectiveness. Despite its significance in matching biotic environments, no TGIP-associated phenotypes were observed for heat-treated offspring. Heat-spikes caused by climate change thus threaten TGIP benefits, potentially increasing susceptibility to emerging marine diseases. This highlights the urgent need to understand how animals will cope with climate-induced changes in microbial assemblages by illustrating the importance – and limits - of TGIP in mitigating the impacts of environmental stressors on offspring vulnerability.

## Introduction

Trans-generational plasticity provides an evolutionary opportunity for short-term acclimatization to changing environmental conditions by shaping the offspring phenotype through the transfer of parental experience (1–4). Trans-generational plasticity can be adaptive and improve offspring survival in situations of rapid but predictable changes in abiotic conditions (e.g., temperature & salinity in marine habitats (5–10)) as well as biotic conditions (e.g., changing microbial assemblage) in a matching parent-offspring environment (4,11,12). In the latter situation, trans-generational immune priming (TGIP; the transfer of parental immunological experience) was suggested to enhance offspring immune responses towards an infectious agent that parents have previously encountered. TGIP, best known from the early discovery of mammalian maternal antibody transfer (13–17), has evolved independently in plants, invertebrates and vertebrates (4,18–23). It encompasses a changed epigenetic programming and/or a direct transfer of parental substances (24), including cells, cytokines and antibodies (25) that together shape neonatal immunity (26,27). Our recently broadened understanding of TGIP mechanisms (4,28–30), in particular, the roles assigned to epigenetic programming as well as the microbiome, has also expanded our understanding of potential routes of TGIP. While traditionally regarded as a maternal trait via deposition into the maternal eggs (19,31) or provisioning through maternal care (pregnancy or post-natal feeding (32–35)), the role of the father in TGIP has recently began to be explored (23,36–43). To this end, TGIP entails a plethora of distinct routes on how immunological experience of the innate or adaptive immune branch can be transferred into the next generation, to support the onset of an effective immunological defense in early life (44–46).

Already a single parental pathogen encounter can result in offspring protection from infectious disease (47) in a matching parent-offspring environment scenario. A mismatch in the microbial assemblage between paternal and offspring environments, as well as a change of an additional abiotic environmental factor across generation, can induce costs of parentally-influenced constitutively upregulated immune responses in the offspring (48,49). While the latter may negatively influence offspring performance, TGIP that relies on inducible immune responses, might become ineffective when additional environmental factors change across generations, due to the resource-allocation trade-off involved (6,50,51). Disentangling constitutive from inducible influences of paternal immunological priming on offspring immune responses can thus illuminate how animals cope in a world of global change where emerging diseases can be a result of rapidly changing abiotic conditions.

The immune system is in close reciprocal interaction with its natural microbiome. The microbiome can strengthen host immune responses, while the immune system shapes microbial niche colonization and composition (52,53). In early life the initial microbial colonization of the gut is crucial for offspring performance and fitness, giving the neonate immune system assemblage and maturation a particularly important role (54,55). The establishment of the gut microbiome can thus be expected to closely interact with TGIP as the latter may potentially directly shape the vertical transfer of microbes supporting gut microbial colonization (56,57). Closing the current knowledge gap of how immune priming shapes the microbiome and, *vice versa*, how the microbiome influences TGIP, may provide insights into the heterogeneity of TGIP within and across populations (58).

While parasites and pathogens are ubiquitous and outrange the diversity and number of hosts in any given habitat (59), the marine realm harbors a particularly high microbial density and diversity (e.g., on average 10_6_ bacteria and >10_7_ viruses / ml of seawater (60). Environmental changes exacerbate the evolutionary mismatch among microbial and eukaryotic interaction partners, characterized by distinct generation times, mutation rates and populations sizes (61). This creates fertile ground for the emergence of marine diseases, often facilitated by rapid shifts in microbial dynamics across the mutualism-parasitism continuum (62). Such scenarios of changing microbial genotype assemblages increase the importance of alternative evolutionary strategies providing a fast acclimatization opportunity to hosts, such as TGIP.

Pipefishes inhabiting coastal seagrass ecosystems are exposed to a high diversity of microbes (63), with *Vibrio* bacteria belonging to both the most abundant bacteria in the water column as well as in the pipefishes’ gut microbiomes (64–68). Over the season, *Vibrio* bacteria may alter from mutualistic members of the microbiome to virulent disease agents through rapid changes in abiotic factors (e.g., temperature and salinity), reinforced by the increasing frequency of heatwaves in recent years (69–71). The pipefishes’ unique male pregnancy makes them prime model systems for the investigation of trans-generational effects (23,37,38,51,72,73). While mothers may transfer their experience directly through the eggs, fathers prime their offspring through their intimate contact during their male pregnancy that evolved with a placenta-like structure in *Syngnathus* and *Hippocampus*, among others (74). Pipefishes and seahorses thus provide the unique opportunity to disentangle the effects of trans-generational immune priming via egg production and pregnancy, that are usually intermingled within the female, by experimentally manipulating the maternal (via the egg) vs. the paternal (during pregnancy) immunological provisioning (4,23,72).

In this study, we have simulated pathogen exposure in female and male broad-nosed pipefish (*Syngnathus typhle*) using two vaccinations with non-indigenous heat-killed *Vibrio* bacteria. We then followed the influence of TGIP on offspring transcriptome-wide gene expression and microbiome composition by experimentally exposing the F1 generation after birth fully-reciprocally to the same strain of *Vibrio* the parents were vaccinated with in two temperature scenarios (18°C “ambient” vs. 25°C “heat-spike”; Fig 1). This design permitted to investigate the following hypotheses: 1) Trans-generational immune priming has a protective effect for the offspring in matching biotic (here: bacterial) parent-offspring environments. 2) TGIP leads to costs associated to induced non-matching phenotypes in non-matching biotic (here: bacterial) environments. 3) Trans-generational immune priming enhances offspring protection specifically against the bacterial strain experienced by the parental generation, while other strains/species remain unaffected. 4) Maternal and paternal TGIP lead to different response patterns in offspring upon bacterial exposure. 5) Strong mismatch in the abiotic parent-offspring environment (here: heat-stress in offspring) can compromise the effectiveness of TGIP.

**Fig 1.**
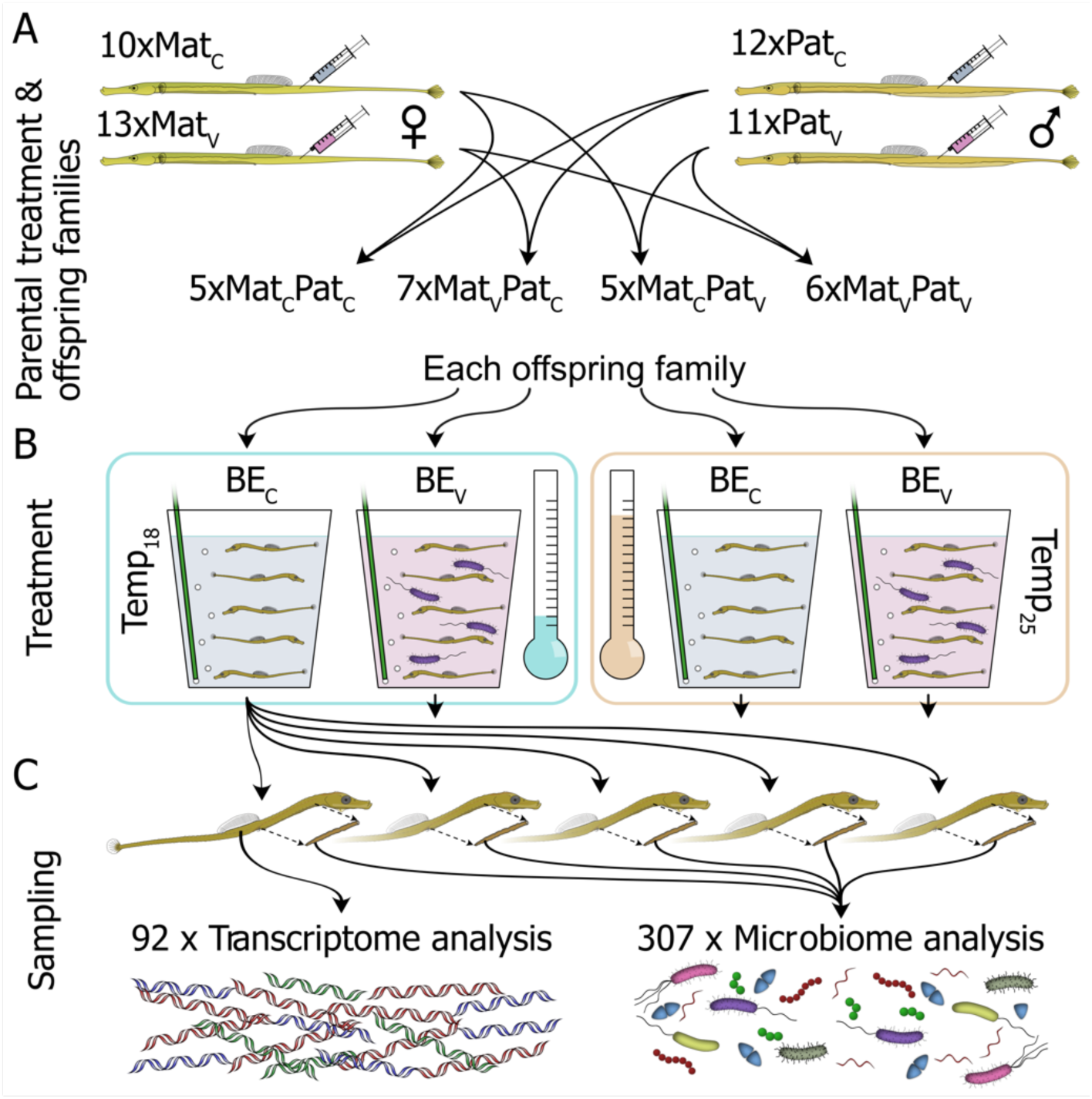
Experimental design for transgenerational immune priming. **(A)** Male and female pipefish were either vaccinated with heat-killed *Vibrio* bacteria (Mat_V_ & Pat_V_) or sham treated with PBS (Mat_C_ & Pat_C_). Parents were mated and 23 juvenile sibling families (with >=20 F1 each) were obtained of which either none, one, or both parents had been vaccinated. (B) Per family, five juveniles were then kept in one of four treatments for one day: seawater with or without *Vibrio*, at 18°C or 25°C. (C) Then, individuals were killed and their guts dissected. From each family treatment group one juvenile without gut was used for RNA extraction and transcriptomics, while the guts of this individual as well as those of other individuals of the same group were used for DNA extraction and microbiome analysis. Given sample numbers are those finally considered throughout analyses.

Our data suggests that TGIP improved offspring defences in a specific manner in matching biotic environments with the parental TGIP effects being distinct and not additive, and costs in mismatching environments appear negligible. The heat-stress treatment reflecting a mis-matched parent-offspring environment scenario revealed how vulnerable this mechanism is as parental effects were virtually absent, probably as a result of a resource-allocation trade-off with a general heat-stress response, emphasizing the threat climate-change induced heat-waves pose to mechanisms natural populations employ to cope with swiftly changing microbial assemblages in their environment.

## Results

### Transcriptome analyses

Gene expression data was preprocessed to remove an unintended effect linked to our tissue dissection procedure as well as to account for the random effect of the fish “family” (Fig 1), yielding gene expression residuals, which were used in all downstream analyses (see Materials & Method & Supporting Protocol). A PCA on these gene expression residuals was computed (Fig 2; Fig S1-3; Supporting Data 1-3) and scores of PC1 to PC4 were tested for differentiation between parental treatment groups (treated as four independent groups, using Dunnett’s test with the control group as reference; package “DescTools “, v. 0.99.47;(75)), temperature groups (Temp_18_ and Temp_25_) and bacterial exposure groups (BE_C_ and BE_V_; for test reports see Supporting Data 3). Scores of immune primed groups (Mat_V_Pat_C_, Mat_C_Pat_V_ & Mat_V_Pat_V_) on PC1-4 did not significantly differ from the control group (Mat_C_Pat_C_; Fig 2A,D,G,J), but scores did significantly differ between Temp_18_ and Temp_25_ on PC1-3 (Fig 2B,E,H) and BE_C_ and BE_V_ for PC 3-4 (Fig 2I,L). BE differences on both PC3 and PC4 thus reflect that some variance linked to the bacterial exposure also negatively covaried with the Temperature treatment (PC3), while some variance does not (PC4). On PC1-2 Temp_18_ and Temp_25_ are significantly different, but the BE groups are not. Interestingly, while Mat_C_Pat_C_ was not statistically significantly different from any other parental vaccination group, score medians of the non-primed control group appear to be lower than in the three primed groups for both PC1 and PC2 (Fig 2A,D). The inverted temperature effect on these PCs (Fig 2B,E) thus suggests that PC1 and PC2 reflect opposing covariance of the temperature and parental vaccination factors, arguing that indeed immune priming is linked to PCA scores, but its effect was split over the first two PCs (and possibly more).

**Fig 2.**
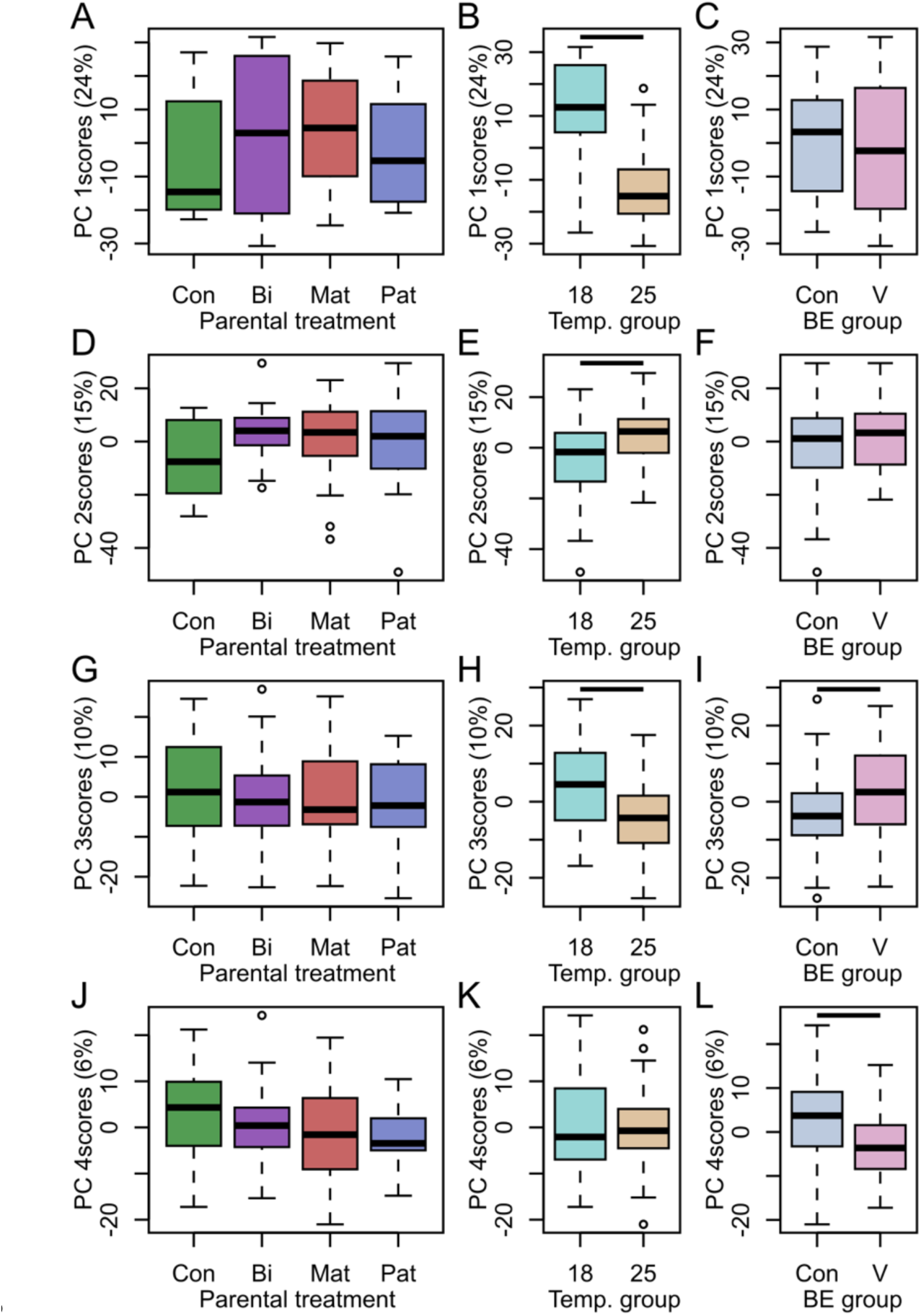
Sample scores and gene loadings from a PCA on preprocessed gene expression data. (A,D,G,J) Sample scores of PC1-PC4, respectively, for parental treatment groups. (B,E,H,K) Sample scores for the Temp treatment groups and (C,F,I,L) scores for the BE treatment groups. Bars indicate significant differences.

To account for the strong and potentially complex Temp effect on gene expression and to unravel the interactions among parental and offspring treatment factors, gene expression differences between selected treatment groups and a reference group were evaluated for each temperature treatment separately using pairwise comparisons based on gene-wise linear mixed models (Fig 3; see Material & Methods and Table S1). Using the group Mat_C_Pat_C_BE_C_Temp_18_ as reference, neither the maternal nor paternal vaccination by itself induced significant changes in offspring gene expression at 18°C after fdr correction when offspring was not exposed to the experimental bacterium (Fig 3A). In offspring from non-vaccinated parents, bacterial exposure (Mat_C_Pat_C_BE_V_Temp_18_) led to 374 differentially expressed genes (DEGs) at 18°C. In contrast, when offspring of a vaccinated mother was exposed to the experimental bacteria, 1,763 DEGs were identified (Mat_V_Pat_C_BE_V_Temp_18_), while only 487 DEGs were found if only the father was vaccinated (Mat_C_Pat_V_BE_V_Temp_18_), and 1,266 genes differed in expression compared to the reference when both parents were vaccinated at 18°C (Mat_V_Pat_V_BE_V_Temp_18_). When the same comparisons were conducted with offspring treated at 25°C (including the reference: Mat_C_Pat_C_BE_C_Temp_25_), bacterial exposure of the offspring (Mat_C_Pat_C_BE_V_Temp_25_) led to only a single DEG. Furthermore, the offspring from vaccinated mothers (Mat_V_Pat_C_BE_V_Temp_25_) showed 80 DEGs after a bacterial exposure at 25°C when compared to the reference - all other considered comparisons did not yield any DEGs. When gene expressions from non-challenged offspring from unvaccinated parents were compared between temperature treatments, 827 differentially expressed genes were identified (Mat_C_Pat_C_BE_C_Temp_18_ vs. Mat_C_Pat_C_BE_C_Temp_25_). Only 27 of these were also among the previously mentioned 374 genes that responded to the bacterial exposure at 18°C (Mat_C_Pat_C_BE_C_Temp_18_ vs. Mat_C_Pat_C_BE_V_Temp_18_), although 26 of these 27 genes showed an expression change in the same direction relative to the reference group (Mat_C_Pat_C_BE_C_Temp_18_ in both cases).

**Fig 3.**
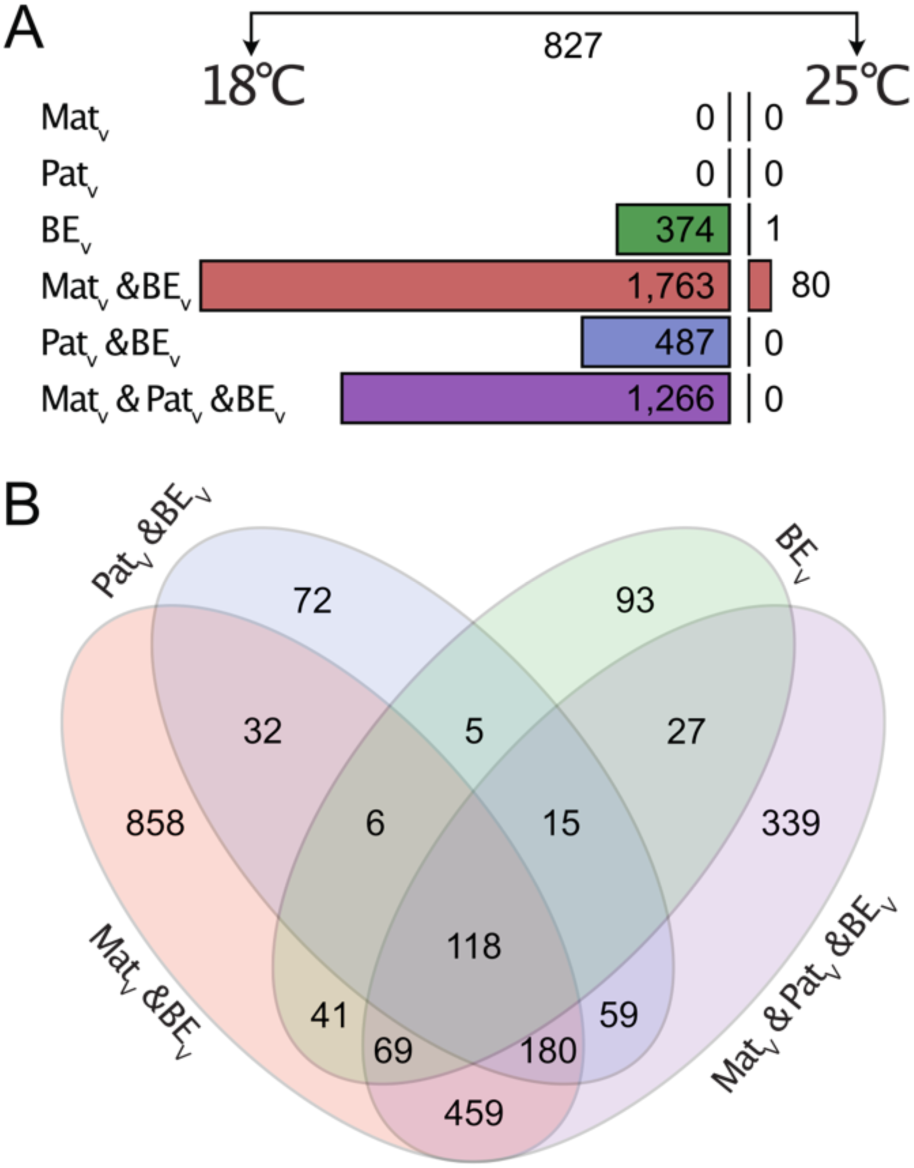
Numbers of differentially expressed genes of pairwise comparisons based on gene expression mixed models. (A) Numbers of significantly differentially expressed genes (after fdr correction) between focal groups and the Mat_C_Pat_C_BE_C_Temp_18_ (left side) or Mat_C_Pat_C_BE_C_Temp_25_ (right side) reference group. (B) VENN diagram of shared differentially expressed genes among comparisons for comparisons with the 18°C reference group.

To evaluate if sets of differentially expressed genes derived from pairwise comparisons were enriched in genes highly expressed in immune cells, we used a custom average immune cell rank analysis (see Materials and Method & Supporting Protocol). For genes differentially expressed between juveniles from non-vaccinated parents without bacterial exposure and juveniles from non-vaccinated parents with bacterial exposure (Mat_C_Pat_C_BE_C_Temp_18_ vs. Mat_C_Pat_C_BE_V_Temp_18_: 374 genes; Fig 4A-D) a significant increase in IM values was detected (p=0.002; suggesting an association with immune cells), as well as significantly higher AD (p=0.032; suggesting an association with cells of the adaptive immune branch), IN (p=0.002; suggesting an association with cells of the innate immune branch) and IA values (p=0.024; suggesting a stronger association with cells of the innate than adaptive immune branch). For the 827 differentially expressed genes between the 18°C and the 25°C control groups (Mat_C_Pat_C_BE_C_Temp_18_ vs. Mat_C_Pat_C_BE_C_Temp_25_) we found on average significantly higher IM values compared to the background (p<0.001; Fig 4E-H), which was also true for AD (p<0.001) and IN (p<0.001) values, but not IA (p>0.05) values, suggesting that there is no bias towards one of the immune branches.

**Fig 4.**
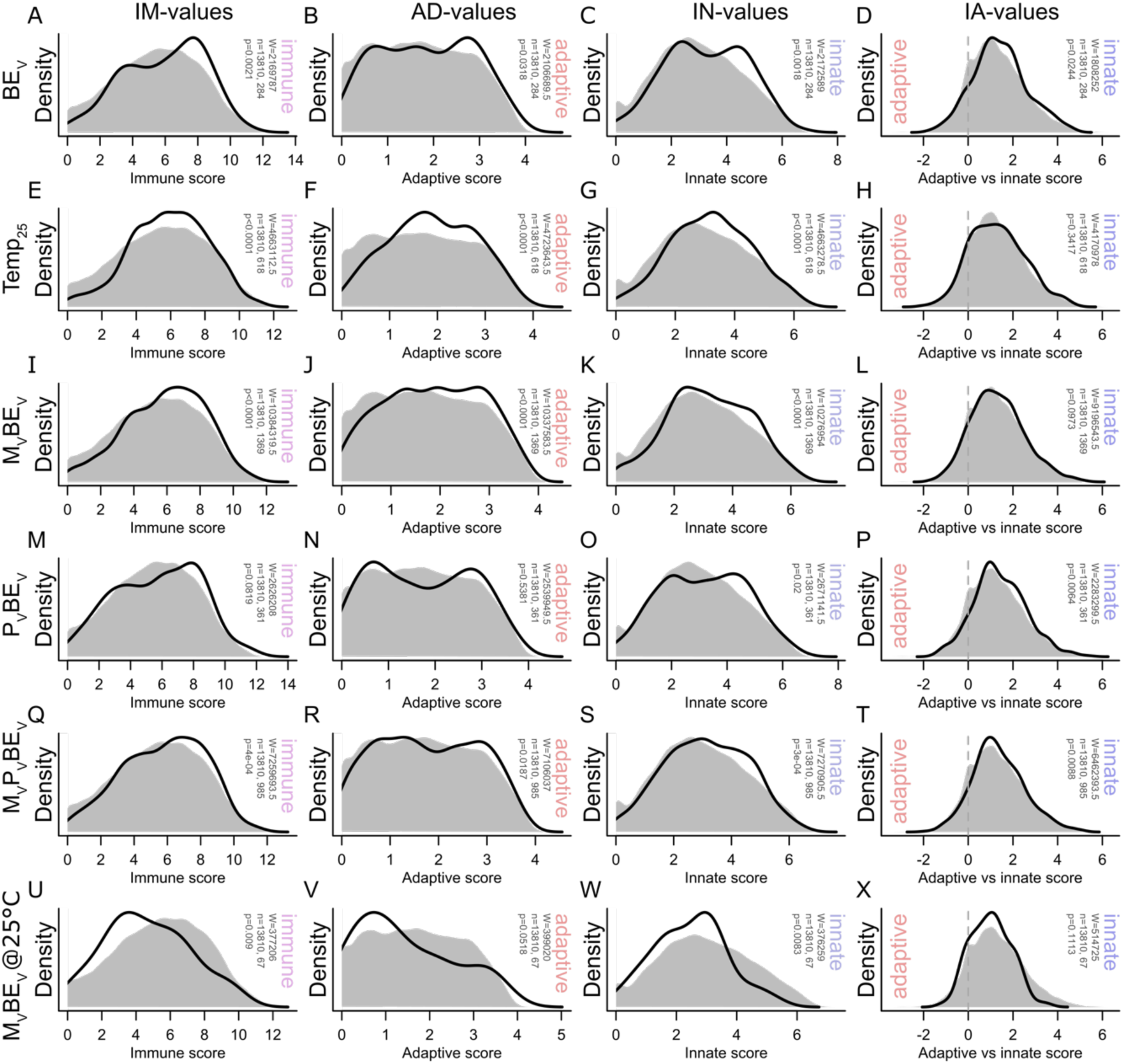
Average immune cell expression rank analyses of target gene sets. (A,E,I,M,Q,U) Distribution of IM-values, (B,F,J,N,R,V) AD-values, (C,G,K,O,S,W) IN-values and (D,H,L,P,T,X) IA-values of significantly differentially expressed genes of selected pairwise comparison shown as black lines (see Fig 3). Reference groups are (A-T) Mat_C_Pat_C_BE_C_Temp_18_ or (U-X) Mat_C_Pat_C_BE_C_Temp_25_. Grey shading represents the distribution of these values among all genes considered in the analysis with human annotation (13,810 genes). Significant differences in median values between sets of significant genes and the background (all genes) were determined using Wilcoxon tests.

Similarly, genes differentially expressed between the bacterial exposure individuals without parental vaccination and those with maternal vaccination (Mat_C_Pat_C_BE_C_Temp_18_ vs. Mat_V_Pat_C_BE_V_Temp_18_: 1,763 genes) also showed on average higher IM, AD and IN scores than the background (all p<0.001; Fig 4I-L), while there was a trend in AI values between this gene list and the background suggesting that possibly this effect is stronger in the innate immune branch than the adaptive one (p=0.097). In contrast, genes differentially expressed between the control group and paternally vaccinated and bacterial exposure individuals (Mat_C_Pat_C_BE_C_Temp_18_ vs. Mat_C_Pat_V_BE_V_Temp_18_: 487 genes) were significantly increased in IN and AI values compared to the background (p=0.02 & p=0.006, respectively; Fig 4O-P), suggesting that differentially expressed genes were indeed specifically linked to cells of the innate immune system. Wilcoxon Signed-Rank tests neither suggested a difference in IM nor AD values relative to the background, but the rather bimodal shapes of the DEGs’ distributions (Fig 4M-N) may differ in variance to the background rather than median – which cannot be detected using our testing regime.

Genes differentially expressed between juveniles without parental vaccination and no bacterial exposure, and those bacterial exposure individuals with biparentally vaccinated parents (Mat_C_Pat_C_BE_C_Temp_18_ vs. Mat_V_Pat_V_BE_V_Temp_18_: 1,266 genes; Fig 4Q-T) showed significantly increased IM, AD, IN and AI values (all p<0.05), suggesting that expression of differentially expressed genes was over-proportionally associated with cells of both branches of the immune system, but also that this effect is significantly stronger for the innate branch compared to the adaptive branch. Finally, the genes that differed at 25°C between the control group and bacterial exposure offspring from vaccinated mothers (Mat_C_Pat_C_BE_C_Temp_25_ vs. Mat_V_Pat_C_BE_V_Temp_25_: 80 genes; Fig 4U-X) showed significantly smaller average IM and IN values (and a trend towards smaller AD values p=0.052) compared to the background, suggesting that expression of these genes was less associated with immune cells than in the background. In summary, we identified an overall trend for differentially expressed genes at 18°C to be associated with genes of the immune system, and especially for the BE treatment and the paternal vaccination groups this affects the two branches of the immune system differently (in both cases the innate branch appears to be more associated to differentially expressed genes), while differentially expressed genes in the maternal vaccination group at 25°C were actually less associated with immune genes than the background.

GO enrichment was explored for genes with significant differential expression in pairwise comparisons (Fig 3A) and only for DEGs of one pairwise comparison enriched terms could be identified (with both human and zebrafish annotations; both annotations can be found in Supporting Data 4: all “pw …” sheets). With human annotation, the six terms enriched in the temperature comparison (i.e., non-bacteria exposed offspring from non-vaccinated parents 18°C vs. 25°C) are linked to catabolic and metabolic processes, but also blood regulation (e.g., “regulation of body fluid levels” and “blood coagulation”). Interestingly, using zebrafish annotations additional GO terms are enriched suggesting immune system involvement: e.g., “complement activation”, “complement activation, alternative pathway”, “killing of cells of another organism” & “disruption of cell in another organism”.

### Microbiome analyses

Complementary to transcriptome analyses, the gut microbiome was genotyped using 16s rRNA barcode sequencing. Obtained barcodes were annotated and a microbiome of 307 samples was computed. Normalized counts of the experimental *Vibrio* strain were extracted per sample, a log(count+1) transformation was applied to them and they were analyzed using a linear mixed model. Model residuals did mostly follow a normal distribution, however, some outliers were identified but tolerated. For the model, log-transformed relative counts were used as dependent variable and Mat, Pat, BE and Temp, as well as all interactions were used as independent variables with the Family variable as random effect and an BE-linked variance structure. As the focus of the analysis was to determine if parental priming induced by parental vaccinations affects microbe counts, the *Vibrio* treatment (and not the control) was chosen as a reference level for the BE factor. The Anova(…, type=”III”) function was used to produce p-values (package “car”). Our model identified significant negative effects for the factors BE, Temp, Mat and Pat (all p<0.05; Table S2; Fig 5). A trend (p=0.076) towards a positive interaction of Mat and Pat might suggest that parental vaccination effects are not additive, but rather redundant. As expected, the most significant and largest effect among these was the BE effect, confirming that treated juveniles showed substantially higher relative abundance of the experimental *Vibrio* strain. A significant interaction of the Pat and Temp factors (p=0.034) with a positive estimate suggests that the negative effect of paternal vaccination on *Vibrio* abundance in guts of offspring experimentally exposed to bacteria was not maintained at higher temperatures and possibly was even inverted. Surprisingly, non-challenged offspring from either mothers or fathers with vaccination appear to have somewhat higher experimental *Vibrio* abundance compared to their control group (the interaction of Pat and BE, and Mat and BE, while not reaching significance, show positive estimates and quite low associated p-values: 0.054 and 0.123, respectively; but Fig 5 illustrates that some outliers may drove this pattern). In contrast, the offspring from biparentally vaccinated parents do not as is reflected in a significant interaction of Mat, Pat and BE (p=0.022) with a negative slope showing that non-challenged juveniles from vaccinated mothers or fathers do have lower experimental *Vibrio* abundance when compared to challenged individuals. A significant interaction of Pat, BE and Temp (p=0.047) with a negative slope illustrates that at 25°C non-challenged paternally vaccinated individuals showed lower experimental *Vibrio* abundance than at 18°C, at which this group appeared to have a higher abundance compared to the BE_C_ control group. The paternal vaccination effect was thus more dependent on environmental temperature than the maternal effect.

**Fig 5.**
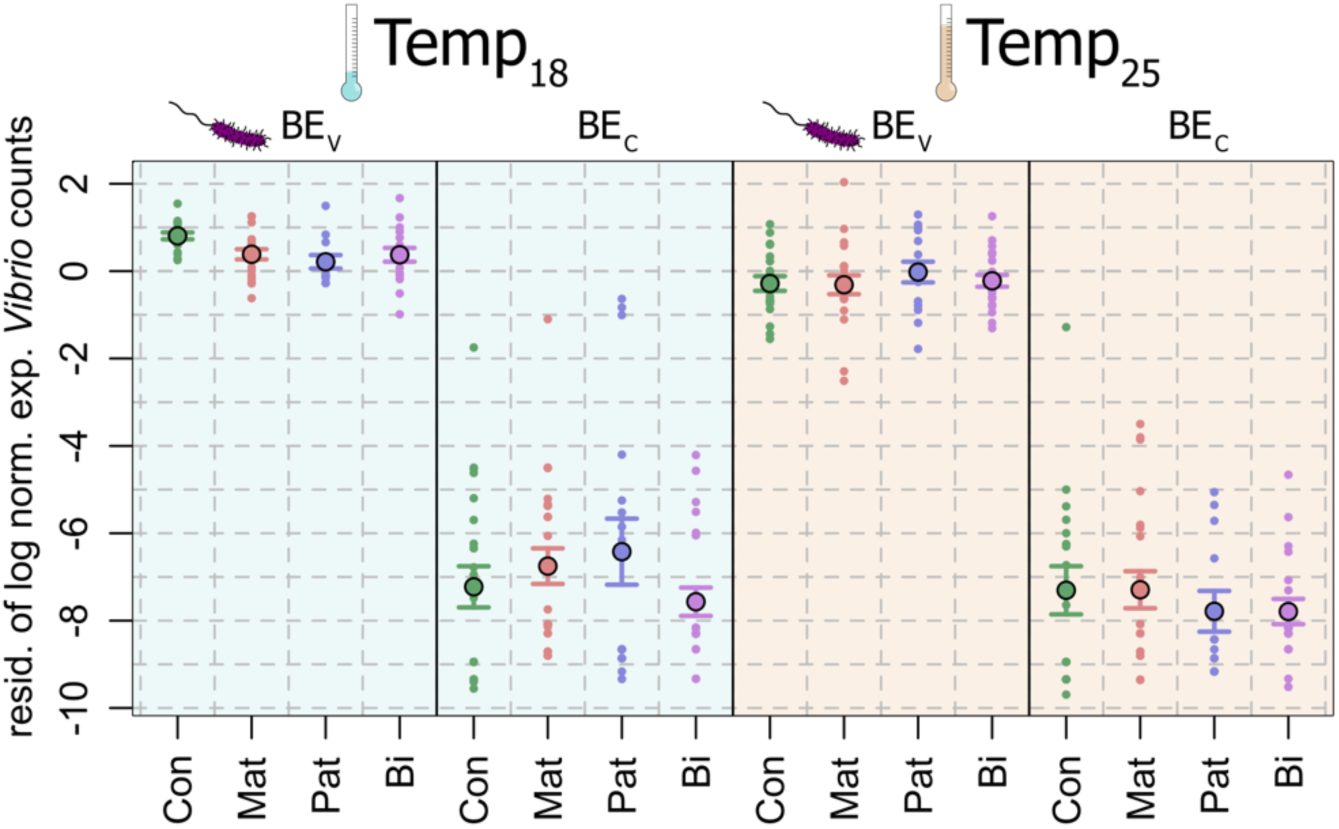
Relative experimental *Vibrio* abundances. Mean and standard error of family-residuals of log-transformed relative experimental *Vibrio* counts across experimental treatment groups.

Experimental *Vibrio* strain relative abundances were then correlated to each gene’s expression values for those individuals of which both data sets were obtained (n=87) and those genes that were also previously used for mixed model analysis (n=16,647). We identified 2,789 genes with Spearman correlation raw p-value smaller than 0.05, however, after adjustment for multiple testing using the false discovery rate procedure only one was considered significant, indicating that this approach for multiple testing might have been overly conservative (as discussed in the Supporting Protocol; Fig S4). We calculated that the number of raw p-values below the selected α of 0.05 exceeded the number of those expected to be observable by chance by 1,957, and thus we selected the 1,957 most significantly correlated genes to analyze their GO term enrichment and immune cell rank composition. GO enrichment was not detected among these genes using a human (or zebrafish) annotation (using a zebrafish annotation, only one term was enriched: “generation of precursor metabolites and energy”). Additionally, the average immune cell expression ranks of these genes were compared to the background ranks (Fig S5). Significantly higher IM, AD and IN values in these genes compared to the background suggest that these genes’ expression levels are on average more associated to cells of the immune system compared to the background, and that both branches of the immune system are equally affected (IA values are not different from the background; p=0.1112).

To determine if parental vaccinations affected gut microbiota compositions beyond the experimental *Vibrio* strain, all other strains’ counts were summed up on the family taxon level per sample and mean count sums were calculated per factor level combination across samples (Fig S6). Most abundant bacteria families excluding the experimental *Vibrio* strain were *Vibrionaceae* (28% of all counts), *Colwelliaceae* (23% of all counts) and *Rhodobacteraceae* (8% of all counts; all others <6%). Based on this family-level table (excl. experimental *Vibrio* counts), the Shannon-Diversity index was calculated per sample, which was found to be negatively correlated to the log abundance of the experimental *Vibrio* strain (Spearman correlation: S= 5495530, rho=-0.14, n=307, p<0.0144), an effect possibly at least partly caused by the reduced read number in bacteria exposed juveniles that were available for non-experimental microbiota strains. A mixed model (after model selection; see Supporting Protocol) also confirmed that the Shannon diversity was negatively correlated with the variable “exp. *Vib*. log(counts+1)” (p=0.0009), while the higher temperature group was associated with higher Shannon diversity values (Table S3). The paternal vaccination effect was also retained in the final model but by itself did not attain significance (p=0.11), however, its interaction with the experimental *Vibrio* count variable (p=0.0087) could suggest that a paternal vaccination effect can affect the Shannon diversity dependent on the prevalence of the experimental *Vibrio* strain: the more experimental *Vibrio*, the more positive the effect of paternal vaccination on alpha diversity. Also, based on the family-level table a significant ANOSIM analysis of offspring exposed to a bacterial exposure at 18°C (R=0.104, p=0.001) revealed that the four parental vaccination groups (Mat_C_Pat_C_, Mat_C_Pat_V_, Mat_V_Pat_C_, Mat_V_Pat_V_; all BE_V_ and Temp_18_) differ in their dissimilarity from the background (although one group (Mat_V_Pat_C_) showed higher average dissimilarity as the background – a pattern we cannot explain; Table S4). In contrast, without bacterial exposure groups did not differ from each other, even with parental vaccination (R=0.0105, p=0.276; BE_C_ and Temp_18_; Table S4).

## Discussion

Adaptive trans-generational plasticity prepares offspring for the specific environmental settings experienced by their parents and it is predicted to increase their survival in a matching environmental abiotic (e.g., temperature, salinity) and/ or biotic (e.g., microbial assemblage) condition across generations. Here, we investigated trans-generational immune priming in the pipefish *Syngnathus typhle* with their unique evolution of male pregnancy, permitting ample opportunities for parent-specific transfer of immunological experience through the maternal (eggs), as well as the paternal (male pregnancy) lineage (23,72). Specifically, we evaluated how trans-generational immune priming upon a parent-specific exposure to heat-killed bacteria affected the transcriptome-wide immunological reaction in offspring. Additionally, we tested, if parental priming affected the bacteria’s abundance in offspring when exposed to the same bacterium, and also how priming and experimental bacteria exposure influenced the offspring’s microbiota. Finally, this was done in a normal vs. a heat-stress environment – an environmental condition expected to be increasingly frequent in the marine realm (76).

Offspring temperature treatment simulating a heat-stress environment of 25°C had a profound effect on gill gene expression (Fig 2B,E,H; Fig 3; Table S1): enriched GO terms were associated with metabolism and catabolism, as well as the complement system (Supporting Data 3 (77)). We also identified several heat shock proteins to respond to our heat treatment (p<0.05), including downregulation of *hspb7*, and upregulation of *hspb1*, *hspa1,* and a trend for *hsp90ab1* (p=0.054; Supporting Data 3), suggesting that the response towards heat stress was metabolically costly (for comparison: no *hsp*-gene changed expression in response to the bacterial exposure (78)). The heat-stress treatment in this study was applied to investigate, how the potentially adaptive effect of TGIP on offspring gene expression and microbial community changes upon non-matching abiotic environmental conditions in the offspring generation (25°C) when compared to ambient temperature 18°C. Therefore, gene expression patterns at 18°C are discussed first.

The offspring exposure with orally supplied *Vibrio* bacteria induced a substantial change in gene expression at the control temperature of 18°C, which becomes apparent when inspecting the PCA results (Fig 2) and numbers of significant interaction estimates across mixed models (Table S1) or pairwise comparisons of gene expression (Fig 3). While GO enrichment was not indicative of any specific group of genes contributing to this response, we found that genes whose expression was negatively correlated with relative experimental *Vibrio* abundance (Fig S5) and DEGs associated with the bacterial exposure (Fig 3A) were significantly associated with immune cells. The latter were significantly more associated with cells of the innate immune system than the background (Fig 4A-D). This suggests that the overall response of *S. typhle* offspring to the experimental *Vibrio* challenge included a modulation of their immune system activity, especially the innate branch, possibly as the adaptive immune system is still subject to maturation, as was suggested for other juvenile teleosts (79–81). Non-exposed juveniles showed significantly lower but still non-zero abundance of the experimental *Vibrio*, which likely reflect naturally occurring closely related *Vibrio* strains present in all individuals that cannot be differentiated from the experimental strain using the V3-V4 region of their 16S rRNA.

Maternal and paternal immune priming affected the offsprinǵ transcriptomic response to exposure with the experimental *Vibrio* bacteria (Fig 4A), which is supported by lower relative *Vibrio* abundance in guts (Table S2; Fig S7), arguing that TGIP-related transcriptomic changes indeed facilitated a stronger or more targeted immune response, which supports our hypothesis 1. However, when offspring was left naïve, no influence of parental immunological exposure on offspring gene expression was identified, suggesting a bacteria-strain-specific effect of the parental *V. aestuarianus* vaccination that only became effective in a matching parent-offspring bacterial environment, i.e., when offspring were exposed to the same bacteria as the parents were vaccinated with, which argues for a generally low cost of TGIP in non-matching biotic (here: bacterial) environments, contradicting our hypothesis 2 (37). Additionally, the lack of a transcriptomic response in primed but not bacterial exposed juveniles argues for a high specificity of the parental immune priming, supporting hypothesis 3. However, parental immune priming seemingly also altered the remaining gut microbial community of juvenile pipefish (Table S3; Table S4): Our ANOSIM suggested that parental immune priming affected microbiota composition. However, our data also showed that the relative abundance of the experimental *Vibrio* affected the composition of the remaining microbiome heterogeneously (Fig S8) and reduced experimental *Vibrio* prevalence (also due to parental immune priming) facilitated a more diverse microbiota in offspring pipefish (Shannon diversity index; higher microbial diversity is generally suggested to be associated with a better health and an improved immune response (82). Thus, we cannot delineate if the microbiome of parentally-primed offspring is different to controls because parental priming also affected non-focal microbial abundances via direct non-specific immune responses, or if these shifts in microbiome composition are merely a result of reduced experimental *Vibrio* abundance due to parental priming. Given that juveniles not exposed to bacteria did not differ in their microbiota composition (despite TGIP), we conclude potential support for hypothesis 3.

The effect of maternal and paternal immune priming differed markedly: relatively few DEGs overlapped between the bacterial exposed maternal and paternal vaccinated groups and a large number of DEGs was associated with the interaction of the parental effects (Table S2; Fig 3B), leading to biparentally vaccinated juveniles not having been significantly different in experimental *Vibrio* abundance from juveniles with unvaccinated parents (p=0.076). This suggest that maternal and paternal TGIP effects are not additive but rather complexly interact, similar to what has been found before in this pipefish species using a candidate gene approach (23) supporting our hypothesis 4. While GO-term enrichment analysis did not provide meaningful insights into how parental effects may differ, our immune-cell association analysis suggests that DEGs of both groups are significantly more associated with immune cells than expected by chance. DEGs of the paternal effect are specifically associated with cells of the innate immune response, while maternal effects are associated equally with both innate and adaptive immune cells (relative to the background; Fig 4I-T). We thus conclude that parental TGIP meaningfully affects pathogen defense in *S. typhle*, but the paternal TGIP contribution involves specifically the innate immune response, which is not the case for the maternal side. Previous studies on candidate genes have found that maternal and paternal priming tended to affect the adaptive and innate immune branch stronger, respectively, which is in line with our results (23,38). The involvement of the innate immune branch in acquired immunity possibly via trained innate immunity could explain how vaccinations function despite the the loss of MHCII and other central immune genes of the adaptive immune branch in *Syngnathus* and other syngnathids (83). We also identified several *hsp*-genes associated with maternal priming and a bacterial exposure (*hsp90aa1*, *hspa4b*, *hspb8*, *hspa1b* all p<0.05), while none was found to be associated with the paternal priming under bacterial exposure. Together with our immune cell association results, this suggests that maternal priming led to a stronger, but more generic stress response, while paternal priming is more specific and more targeted via a modulation of the innate immune branch, possibly suggesting that “trained immunity” may play a pivotal role in syngnathids’ immune biology, as suggested before (84). However, the analysis of experimental *Vibrio* prevalence suggests no significant difference in maternal vs. paternal vs. biparental immune priming effects. The involvement of different immunological pathways in the offspring upon maternal vs. paternal immune priming thus seems to facilitate a similar protective effect when it comes to an exposure of the bacteria previously encountered by the parental generation. The cost of TGIP in the non-matching parent-offspring microbial environment scenario seems low, given that neither maternal nor paternal immune priming led to detectable transcriptomic changes in the offspring generation if left immunologically naive (non-matching biotic condition; Fig 3A), which contradicts hypothesis 2. Parental immune priming in the pipefish *Syngnathus typhle* against *Vibrio aestuarianus* is probably a highly specific and thus likely cost-effective mechanism to boost offspring immune defenses in matching parent-offspring environments. However, studies showing similar TGIP patterns using more varied microbes (for vaccination & offspring exposure) are necessary to truly shed light on TGIP specificity patterns in *S. typhle*.

TGIP patterns in matching biotic parent-offspring environments were virtually absent in the heat-stress treatment (non-matching abiotic parent-offspring situation). On the transcriptome level, the increased temperature led to a substantial shift in gene expression (Fig 2-3; Table S2) and associated DEGs were significantly associated with immune cells of both immune system branches. However, at 25°C only a single gene was identified to respond to the offspring bacterial exposure treatment, expression changes of only 80 genes could be linked to maternal priming when individuals were also exposed to bacteria, and expression of not a single gene could be linked to paternal or biparental priming when offspring was challenged (Fig 3). Furthermore, maternal priming DEGs were significantly less associated to immune cells than expected by chance, indicating that the maintained response may not be immune-related. Three scenarios might have led to this pattern: (i.) the immune response (incl. immune priming) might have been substantially less pronounced at heat-stress conditions. As transcriptomic responses are likely linked to energetic costs, responding to external stressors would presents a resource allocation trade-off scenario supporting hypothesis 5. Under severe heat-stress, as applied in this experiment, available energy might have predominantly been allocated to the heat-response to maintain metabolic homeostasis, hampering the protective effect of TGIP to the bacteria exposure (85). In our experiment, juveniles were only treated for one day and mid-to long-term effects of possibly reduced immunological vigilance (in e.g., survivability, growth rates) could unfortunately not be observed. (ii.) Alternatively, the less pronounced immune response could have been sufficient as fewer *Vibrio* bacteria successfully infected the guts of the heat-stress treated offspring, possibly because environmental conditions were less favorable for an infection with the experimental *Vibrio* strain (relative to other microbes) at high temperatures (25°C) when compared to the 18°C temperature treatment (Table 2; Fig 5 (86)). However, initial experimental *Vibrio aestuarianus* doses were identical between temperature treatments, and *V. aesturianus* growth in *in vitro* cultures at 25°C outperforms growth at 18 °C (results not shown) entailing that the lower *Vibrio* prevalence in offspring gut identified at 25°C is unlikely to be caused by reduced *Vibrio* growth in pipefish gut upon exposure. The latter would also be unlikely given the short time the offspring were observed after the *Vibrio* exposure. (iii.) Finally, the transcriptomic responses to heat-stress and pathogens may overlap to some extend in juvenile *S. typhle*: our analysis suggests that genes responding to heat-stress were over-proportionally associated with immune cells and thus this broad stress response may have already included most genes of a bacterium-induced response. Such an effect might also explain why experimental *Vibrio* prevalence overall was lower in the heat treatment compared to the normal temperature group as was shown by a recent study in which 90% of the genes responding to a *Vibrio alginolyticus* bacterial exposure in Herring (*Clupea harengus*) larvae had also responded in the same direction to a heat stress treatment (87). However, in the present study only 27 DEGs overlap between the pairwise comparisons of Mat_C_Pat_C_BE_C_Temp_18_ vs. Mat_C_Pat_C_BE_C_Temp_25_ (827 DEGs) and Mat_C_Pat_C_BE_C_Temp_18_ vs. Mat_C_Pat_C_BE_V_Temp_18_ (374 DEGs; Supporting Data 2).

While 26 of these 27 DEGs showed indeed an expression change in the same direction in response to the stressor, the overall small number of shared DEGs does not support the hypothesis that the heat-stress and pathogen-stress response has induced similar gene expression patterns in *S. typhle* offspring. Instead, we believe that overall reduced relative experimental *Vibrio* prevalence is likely a result of increased abundance of other microbes that benefit more from the increased water temperature, potentially leading to a relatively smaller number of experimental *Vibrio* counts in the sequenced DNA. We therefore conclude that the first explanation (i.) is the most likely, which raises concerns on fish larval immunological vigilance under heat-stress scenarios that are expected to increase in frequency and severity over the next decades (76). Our data suggests that heat stress does not only lead to a severe change in transcriptomic profiles, but also that immunological vigilance can be compromised likely due to a resource allocation trade-off, as has been discussed before also for other environmental stressors and which supports hypothesis 5 (88,89). Thus, increases in environmental temperature regimes may threaten this pipefish’s natural populations as efficient immune priming may not be possible anymore hampering the resilience towards relatively sudden shifts in microbial compositions expected in their environment.

For host organisms, transgenerational immune priming can mediate the challenges of an environmental microbiome composition that changes too fast for genetic adaptations but slow enough for parent-offspring environmental matching to occur. Vaccinations with heat-killed bacteria in parents, simulating pathogen exposure, can induce effective and specific immune priming in offspring, with mothers and fathers affecting the offspring’ immune responses differently, protecting from pathogens already encountered in the parental generation. This adaptive trans-generational plasticity seems highly cost-effective in a situation of non-matching biotic environments given that the transcriptomic profile of juveniles not exposed to the focal bacterium is not different from a non-primed individual. However, natural heatwave scenarios, which are expected to become more frequent due to on-going environmental change, have the potential to inhibit the positive effects of trans-generational immune priming almost entirely, possibly *via* a resource-allocation trade-off. This raises substantial concerns on how animal will cope with climate change-induced shifts in pathogen assemblages, ultimately predicting alarming scenarios of emerging marine disease.

## Material and Methods

### Experimental design and fish sampling

*Syngnathus typhle* individuals were caught in the seagrass meadow in Falckenstein (54.394497, 10.188988) in April 2020 (*prior* mating season), brought to the Helmholtz Centre for Ocean Research Kiel (GEOMAR) and kept in sex-specific 200L barrels with Baltic Sea water flow through. The temperature was gradually increased from 10°C to 18°C and a day:night rhythm of 16h:8h was achieved over three weeks to approximate summer conditions in the Baltic Sea, inducing fish’s mating readiness. Animals were transferred to 80l aquaria attached to a common water circulation system and kept in same-sex groups (Fig 1A). To induce immune priming in offspring, first, half of the males and females were vaccinated subcutaneously with 50 µl heat-killed bacterial cells (10^8^ *Vibrio aestuarianus* bacteria/ml strain “I11Ma2”; originally isolated from a pipefish foregut caught in the Mediterranean Sea, Italy (90)) twice, with one week in between injections. One day after the second vaccination, eight mating pairs (“families”) were established for each of the intended parental mating groups in separate tanks attached to the same water flow-through system (32 pairs in total): neither female (maternal treatment; “Mat”) nor male vaccinated (paternal treatment; “Pat”; group Mat_C_Pat_C_), only female vaccinated (Mat_V_Pat_C_), only male vaccinated (Mat_C_Pat_V_), and both female and male vaccinated (Mat_V_Pat_V_). The pairs were given two days for mating, after which females were removed, resulting in 23 families with sufficient offspring numbers after male pregnancy for the following experiments (final family numbers were Mat_C_Pat_C_: 5, Mat_V_Pat_C_: 7, Mat_C_Pat_V_: 5, Mat_V_Pat_V_: 6). Body length and weight were recorded for all parents.

Immediately after parturition, the offspring was transferred to a 1l tank per parental tank in a flow through system at 18°C, with the same day and night cycle as before and fed twice a day with live *Artemia salina* nauplii (Fig 1A). Nine to ten days after birth, 20 offspring individuals per family were randomly selected for further treatments and, as offspring of different families varied in age (matings and parturitions were not perfectly synchronized across mating pairs), subsequent experiments were performed on three consecutive days to match offspring age. As family was treated as a random factor in our statistical analyses, a potential effect of the sampling day is also accounted for as all individuals per family were processed on the same day. The 20 individuals per family were split into four groups of equal sizes and each group was moved into a beaker with 300 ml Baltic seawater, which was aerated by a weak air bubbler (Fig 1B). Two beakers per family were kept at 18°C (control temperature; “Temp_18_”), while the other two were kept at 25°C (heat-stress situation; “Temp_25_”; temperature treatment = “Temp”). Finally, juveniles were subjected to a bacterial exposure (“BE”): 50µl of a solution containing live *Vibrio* (again strain I11M2; 10^8^ *Vibrio* bacteria/ml) was added to 50ml rinsed *Artemia* nauplii solution (1dph). After an incubation period of 30min, one milliliter of this mixture was added always to one of the two beakers per family and temperature (“BE_V_”), while the other received the same amount of non-enriched *Artemia* nauplii (“BE_C_”). After one day in the beakers, all juveniles were killed via a lethal dosage of MS222 (0.04% in PBS), an overview photo was taken of all 5 juveniles per beaker to estimate treatment batch average body lengths using the segmented line tool in ImageJ (v.1.52k), and juveniles were moved to RNAlater, kept for 1d at 4°C and then stored at -20°C. Unfortunately, keeping and treating five juveniles within a single beaker induced a nesting factor we cannot correct for (affecting the microbiota dataset exclusively), however, individual-specific beakers were not feasible considering the large sample size. We therefore continued the analysis as if this effect would not exist, but acknowledge that conclusions from the microbiota dataset have to be drawn cautiously.

### Gut dissections

To investigate the gut microbiome of juveniles, a transversal cut through the neck of each RNAlater-stored specimen was conducted, still leaving the ventral body half connected to the trunk. Using forceps, the head (and attached tissues) was then pulled from the trunk, which led the ventral body wall to rip-off directly posterior to the head, while the intestines remained connected to the head and instead got dislodged from the trunk close to the vent. Subsequently, the majority of the intestine was dissected from the head part, leaving only the most anterior part of the gut and neighboring organs, such as the liver, with the head. Intestines were stored in 100% EtOH for microbiome analyses, while all remaining tissues were immediately used for RNA extraction.

### RNA extraction, library synthesis and sequencing

Total RNA was extracted from one individual per family and treatment batch (i.e., “beaker”), leading to a total of 92 samples (23 families, with one individual for each of the two Temp and BE groups). For this purpose, remaining juvenile tissues (i.e., everything but intestines) were taken immediately after gut dissection to homogenization in lysis buffer and processed using the Qiagen DNA/RNA All-Prep Blood and Tissue Mini kit (Cat. No. 80284). Extracted RNA had an average concentration of 41ng/µl and an average RIN of 9.7. mRNA library synthesis and sequencing (DNBseq, 150bp PE, stranded) was conducted by BGI (HongKong, China) and on average 58mio reads/sample were obtained, with all samples having >30mio reads, despite some duplication levels were determined to be rather high, which might be reflected in some data noise.

### Transcriptome assembly

The RNA reads were quality trimmed with Fastp (v0.20.1 (91)). Alignments to the annotated genome were carried out with STAR (v2.7.9a (92)) in a two-pass mode with the following options “--outFilterIntronMotifs RemoveNoncanonical --outSAMunmapped None --outFilterMultimapNmax 1”. The count information per sample was obtained using TPMCalculator with options “-c 150 -p -q 255 -e -a” and then merged into a single multi-sample file with a script provided by the developer (tpmcalculator2matrixes.py). The output files from TPMcalculator were processed using a custom python script that for each *S. typhle* gene added an orthologous gene ID and gene description from *Danio rerio*, *Hippocampus erectus*, *Syngnathus acus* and *Homo sapiens*. Missing orthology information was marked as NA. The orthology assignment was performed with Orthofinder (v2.4.0 (93)) using protein sequences from the following species: *Latimeria chalumnae*, *Danio rerio*, *Xenopus tropicalis*, *Hippocampus comes*, *Hippocampus erectus*, *Syngnathus acus*, *Syngnathus typhle* (this annotation), *Homo sapiens*, *Mus musculus*. 26,072 genes were processed and orthology was identified for *H. sapiens* in 14,435 genes, for *D. rerio* in 16,383 genes, for *S. acus* in 18,197 genes and for *H. erectus* in 19,055. TPM tables were imported into R and a pre-analysis suggested that the applied dissection method affected gene quantification and details are found in the Supplementary Protocol (94). Briefly, variations in sample-wise TPM histograms, a PCA and GO term enrichment analysis suggested that the major axis of gene expression variation was not linked to any intended treatment, but instead linked to differences in sample dissection. The effect was addressed per gene by fitting a linear regression model using PC1 of the afore mentioned PCA as independent variable, log-transformed expression values as dependent variable (log(expr.+1)), and then by only considering the residuals of this model in downstream analyses (see below; Supporting Data 1: sheet “tpm PC1 residuals”).

Gene residuals were used as dependent variables to compute gene-wise linear mixed models, with treatments and their interaction as independent variables and allowing for random intercepts for the “family” factor. Based on these models, pairwise-comparisons between always one of seven treatment groups (Mat_V_Pat_C_BE_C_Temp_18_, Mat_C_Pat_V_BE_C_Temp_18_, Mat_C_Pat_C_BE_V_Temp_18_, Mat_C_Pat_C_BE_C_Temp_25_, Mat_V_Pat_C_BE_V_Temp_18_, Mat_C_Pat_V_BE_V_Temp_18_ and Mat_V_Pat_V_BE_V_Temp_18_) and a reference group (Mat_C_Pat_C_BE_C_Temp_18_: Supporting Data 2: “pairwise 18°C” sheets, or Mat_C_Pat_C_BE_C_Temp_25_: Supporting Data 2: “pairwise 25°C” sheets) were computed and resulting p-values corrected for multiple testing using the “fdr” method. Additionally, as potential family effects would also affect a PCA ordination, another similar linear mixed model was calculated per gene, using no independent variables and only a random intercept to obtain residuals for a family-effect-corrected PCA, whose axes’ scores were tested for difference using Dunnett’s Test (more details in the Supplementary Protocol). For each comparison that yielded genes with fdr corrected p-values below α=0.05, Gene Ontology (GO) terms were assigned to all genes when available and tested for enrichment as described in the Supplementary Protocol (Supplementary Protocol; Supporting Data 3).

### Average immune cell expression rank analyses

To understand if DEGs derived from pairwise-comparisons may have an association with immune functions, we tested DEG lists for enrichment in genes with previously published pronounced expression in cells of the immune cells, cells of the innate-or adaptive immune system, via three indicator scores “IM”, “IN” and “AD”, respectively, with each DEG receiving one value each, as well as all other genes to produce a reference distribution (see Supporting Protocol for details). We also subtracted AD scores from the IN scores per gene (resulting in the “IA” score) to indicate if DEGs were more associated to cells of one immune branch over the other. Generally, higher values indicate a stronger association of genes with cells of the certain type and enrichment was evaluated per score and DEG list by comparing the score distribution median to the background distribution median via Wilcoxen Signed-Rank tests. For these analyses it should be kept in mind that gene sets are not independent of each other, many genes overlap among them and p-values derived from Wilcoxen Sign-Rank tests were not corrected. Also, not all genes had a human annotation and thus actual sample sizes for comparisons were somewhat smaller than DEG lists suggest.

### Microbiome sequencing, assembly and analysis

317 gut samples were processed using the Qiagen DNA Blood and Tissue kit (Cat. No.: 69504). Obtained DNA was sent to the IKMB sequencing center, Kiel, Germany, which performed library preparation and Illumina MiSeq amplicon paired-end sequencing of the V3-V4 bacterial 16S rRNA region randomly distributed to three runs (run 54: n=30, 55: n=8 and 57: n=269). Sequences were demultiplexed, conjoined with their respective metadata and processed, quality-filtered and analyzed (package “QIIME2”, v.2021.8 (95)). Correction of the fastq files was achieved using the DADA2 software within the QIIME2 pipeline, which included denoising, merging, chimera slayer fitting and trimming of the sequences (“ASVs”). Due to declining quality scores, sequences were truncated 50 bases from forward and 70 bases (run 54) or 100 bases (run 55, run 57) respectively from reverse reads. Taxonomy was assigned using a Naïve Bayes classifier trained on SILVA138 99% OTUs full-length sequence database. Microbiota count and meta data of all 317 samples were imported into R using the function qza_to_phyloseq() (package “qiime2R” v.0.99.6 (96); 89 samples of these were also used for gene expression analysis), with reads being assigned to 7,100 microbial taxa (ASVs). Samples with log(counts+1)<=8 (∼3000 reads) were excluded after histogram inspection (leaving 307 samples; minimum sample size per subgroup was n=13 and maximum number n=23) and remaining samples’ counts were normalized by dividing them by the respective sample’s mean count across taxa, and then multiplying them by the mean count across all samples and taxa. Thus, when the text refers to “counts” for brevity, “normalized counts” or “relative abundance” is actually meant subsequently.

Mean log(count+1) of all *Vibrio* ASVs were visualized using a heatmap (Fig S7). One strain (formerly two ASVs of identical barcode sequences were merged into one) showed not only the highest overall abundance among ASVs (16.5% of all microbiome counts), but also featured a marked difference in abundance between bacterial exposure groups indicating that it represents the experimental strain. The experimental strains abundance across samples that also had RNA-seq data (n=87) did not correlate with the bias-indicating original RNA-seq PCA PC1 (see Supporting Protocol), suggesting that no correction was required (S = 127812, p-value = 0.1273). Treatment effects on the experimental *Vibrio* counts were analyzed by first log-transforming all 307 samples (log(counts+1)), followed by visual inspection across treatment groups nine outliers were excluded. Remaining samples’ transformed *Vibrio* counts were used as the dependent variable in a linear mixed model (lme() function in R, with maximum likelihood method). A qq-plot on residuals was used to investigate normality of residuals and while some deviation from normally distributed residuals was noted, overall model validation indicated a robust model. As in the gene expression analyses, independent variables were the factors Mat, Pat, BE and Temp, and all their interactions. A random intercept for the family factor was included, but here also a variance structure for the BE-treatment. The Anova() function was used again to obtain an ANOVA table and type III ANOVA was chosen as interactions rather than main effects were of central interest. The experimental strain counts were also correlated to each gene’s expression value using Spearman rank correlations for those samples of which both microbial count data and gene expression data was obtained. Genes deemed significantly correlated to the experimental *Vibrio* count (see Supporting Protocol) were analyzed for GO enrichment and their average immune cell expression rank, as detailed in the supplemental protocol.

For analyses of the remaining microbiome, first, counts of the experimental *Vibrio* strain were removed and samples re-standardized. Then, all counts from within each microbial family taxon were summed up per individual and families with a total of less than ten counts were removed (332 microbial families remaining). To test if the immune-challenge phenotypes elicited by parental immune priming also affected this remaining microbiome, two ANOSIM analyses were conducted using Bray-Curtis dissimilarity matrices: one on microbiome abundances from samples without bacterial challenge at 18°C (Mat_C/V_Pat_C/V_BE_C_Temp_18_), and one on samples with this exposure (Mat_C/V_Pat_C/V_BE_V_Temp_18_; anosim(); “vegan” package v.2.6-2), for both of which the factor level combinations of Mat and Pat were treated as one factor with four levels. The Shannon Diversity index was calculated per individual using log(counts+1) and its correlation with experimental *Vibrio* counts assessed using a Spearman rank correlation. Additionally, two linear mixed models were computed using the alpha diversity as response variable, and Mat, Pat, Temp and either BE or the experimental *Vibrio* log(counts+1), as well as all interactions as predictor variables (“family” was added as a random effect). For both models model selection using the stepABE() function was conducted and while both models yielded similar BIC values (200.97 and 199.49, respectively), *Vibrio* counts were suggested as a slightly better predictor than BE and was therefore evaluated using the Anova(…,type=”III”) function.

## Supporting information

Supporting Protocol

## Acknowledgements

We would like to thank Kim-Sara Wagner for assistance with the microbiome analysis and Johannes Hasse & Fabian Wendt for support with the animal care. We are grateful to the MarEvol Group at CAU Kiel for their support while experiments were conducted. We thank Thorsten Reusch for hosting our group at GEOMAR Helmholtz Centre for Ocean Research until 2021. We thank the Collaborative Research Center 1182 “Origin and Function of Metaorganisms” and the Institute of Clinical Molecularbiology (IKMB) Kiel for 16s rRNA genotyping.

## Data Accessibility and Benefit-Sharing

Raw reads of the transcriptomic and microbiome datasets can be supplied upon request and accession numbers will be available in the article published after peer-review. All processed data can be found in the Supporting Files.

## Author Contributions

OR and RFS conceived the experiment. OR and SMM conducted the empirical experiments with support by RFS. AD conducted the transcriptome assembly and RFS conducted the statistical analyses, supported by AD and OR. OR and RFS wrote the initial draft of the manuscript and all authors revised and approved it.

## Grant Sponsor

European Research Council (ERC) to Olivia Roth (MALEPREG: eu-repo/grantAgreement/EC/H2020/755659).

DFG via the Research Training Group for Translational Evolutionary Research (RTG 2501 TransEvo)

## Ethics approval statement

All experiments were conducted in accordance with local ethics regulations (University of Kiel Antrag §7 V242-35168/2018.

## Supplemental Tables

**Table S1:** Numbers of significant p-value of the gene expression mixed model estimates. Shown are variables/interactions that had at least one significant fdr p-value or one p-value expected to be significant (pcor=yes; see Supplemental Protocol).

| Variable/interaction | raw p | fdr p | $p_{cor}=yes$ |
| --- | --- | --- | --- |
| Matv | 2,389 | 2 | 1,556 |
| Patv | 881 | 0 | 48 |
| BEv | 5,460 | 3,116 | 4,627 |
| Temp25 | 10,784 | 10,243 | 9,951 |
| Matv:Patv | 1,859 | 3 | 1,026 |
| Matv:BEv | 1,344 | 0 | 511 |
| Matv:Temp25 | 1,310 | 1 | 477 |
| BEv:Temp25 | 5,859 | 3,924 | 5,026 |
| Matv:Patv:BEv | 1,669 | 0 | 836 |
| Matv:BEv:Temp25 | 1,110 | 0 | 277 |
| Matv:Patv:BEv:Temp25 | 1,020 | 0 | 187 |

**Table S2:** Statistical summary of the linear mixed effects model with experimental *Vibrio* log(count +1) as dependent variable and Mat, Pat, BE and Temp, as well as all interactions as independent variables

| Variable | estimate | $\chi^2$ | Df | $Pr(>\chi^2)$ | |
| --- | --- | --- | --- | --- | --- |
| (Intercept) | 10.24 | 1376.71 | 1 | <0.0001 | *** |
| Matv | -1.14 | 9.13 | 1 | 0.003 | ** |
| Pat<sub>v</sub> | -1.02 | 5.87 | 1 | 0.015 | * |
| BE<sub>c</sub> | -8.1 | 318.67 | 1 | <0.0001 | *** |
| Temp<sub>25</sub> | -1.14 | 22.76 | 1 | <0.0001 | *** |
| Mat<sub>v</sub>:Pat<sub>v</sub> | 1 | 3.15 | 1 | 0.076 |  |
| Mat<sub>v</sub>:BE<sub>c</sub> | 0.97 | 2.38 | 1 | 0.123 |  |
| Pat<sub>v</sub>:BE<sub>c</sub> | 1.34 | 3.72 | 1 | 0.054 |  |
| Mat<sub>v</sub>:Temp<sub>25</sub> | 0.45 | 1.84 | 1 | 0.175 |  |
| Pat<sub>v</sub>:Temp<sub>25</sub> | 0.8 | 4.47 | 1 | 0.034 | * |
| BE<sub>c</sub>:Temp<sub>25</sub> | 1.08 | 2.58 | 1 | 0.108 |  |
| Mat<sub>v</sub>:Pat<sub>v</sub>:BE<sub>c</sub> | -2.15 | 5.28 | 1 | 0.022 | * |
| Mat<sub>v</sub>:Pat<sub>v</sub>:Temp<sub>25</sub> | -0.71 | 2.01 | 1 | 0.156 |  |
| Mat<sub>v</sub>:BE<sub>c</sub>:Temp<sub>25</sub> | -0.94 | 1.02 | 1 | 0.312 |  |
| Pat<sub>v</sub>:BE<sub>c</sub>:Temp<sub>25</sub> | -2.08 | 3.96 | 1 | 0.047 | * |
| Mat<sub>v</sub>:Pat<sub>v</sub>:BE<sub>c</sub>:Temp<sub>25</sub> | 2.31 | 2.79 | 1 | 0.095 |  |

**Table S3:**
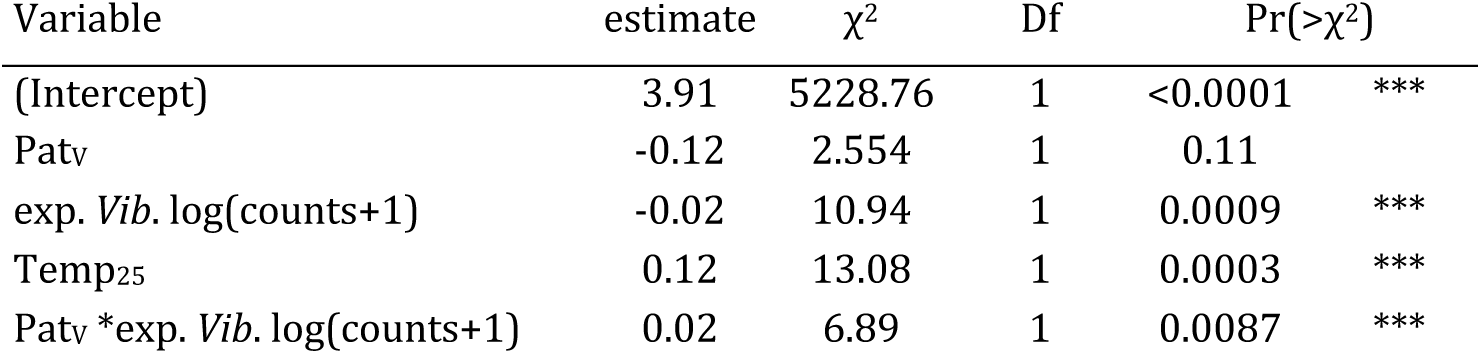
Statistical summary of the selected linear mixed effects model analyzing microbiome diversity. Full model included the Shannon diversity as dependent variable and Mat, Pat, exp. *Vibrio* log(count +1) and Temp, as well as all interactions as independent variables and Fam as random effect.

| Variable | estimate | $\chi^2$ | Df | Pr(>$\chi^2$) |
| --- | --- | --- | --- | --- |
| (Intercept) | 3.91 | 5228.76 | 1 | <0.0001 *** |
| Pat<sub>v</sub> | -0.12 | 2.554 | 1 | 0.11 |
| exp. *Vib.* log(counts+1) | -0.02 | 10.94 | 1 | 0.0009 *** |
| Temp<sub>25</sub> | 0.12 | 13.08 | 1 | 0.0003 *** |
| Pat<sub>v</sub> * exp. *Vib.* log(counts+1) | 0.02 | 6.89 | 1 | 0.0087 *** |

**Table S4:**
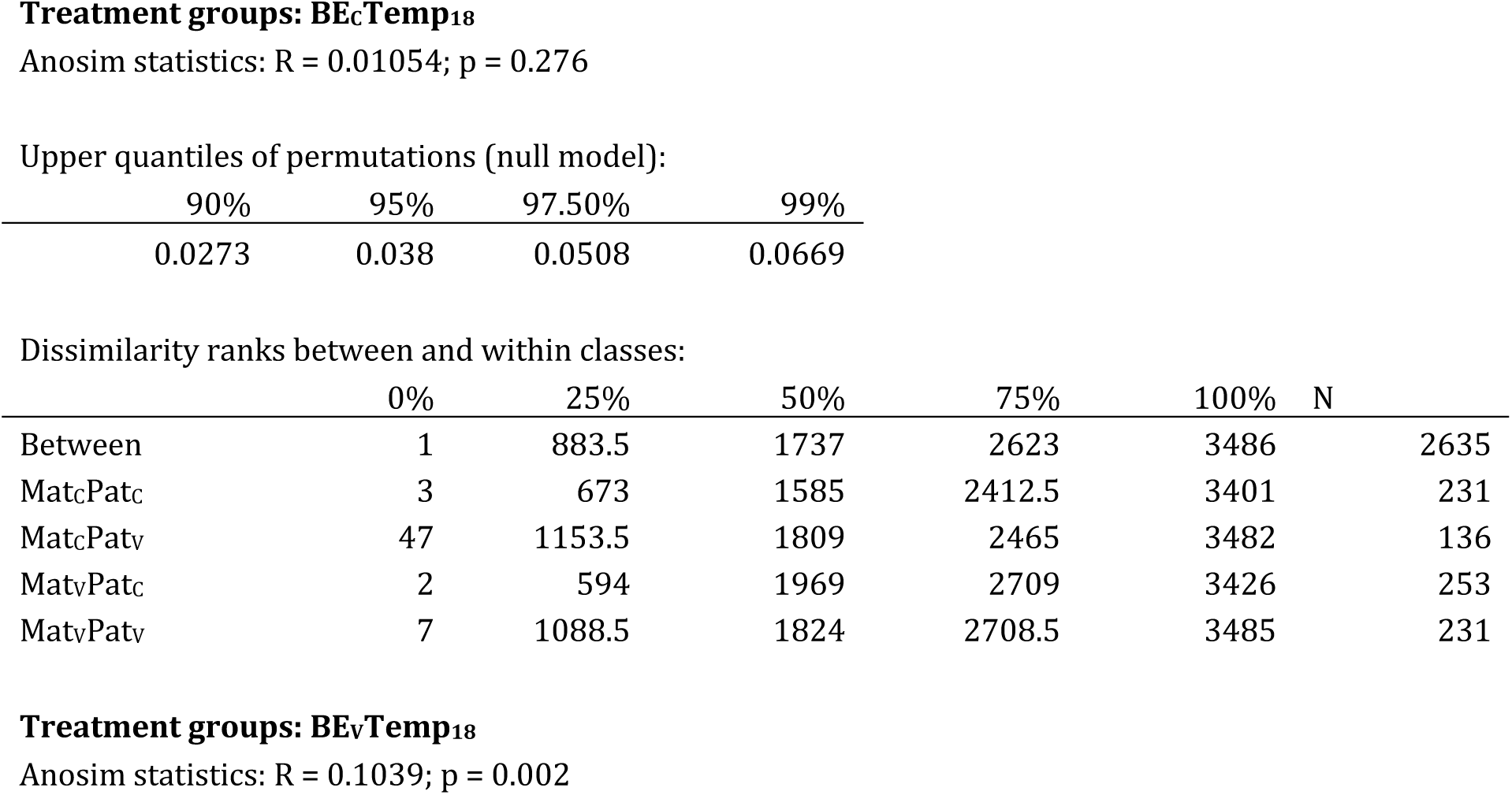
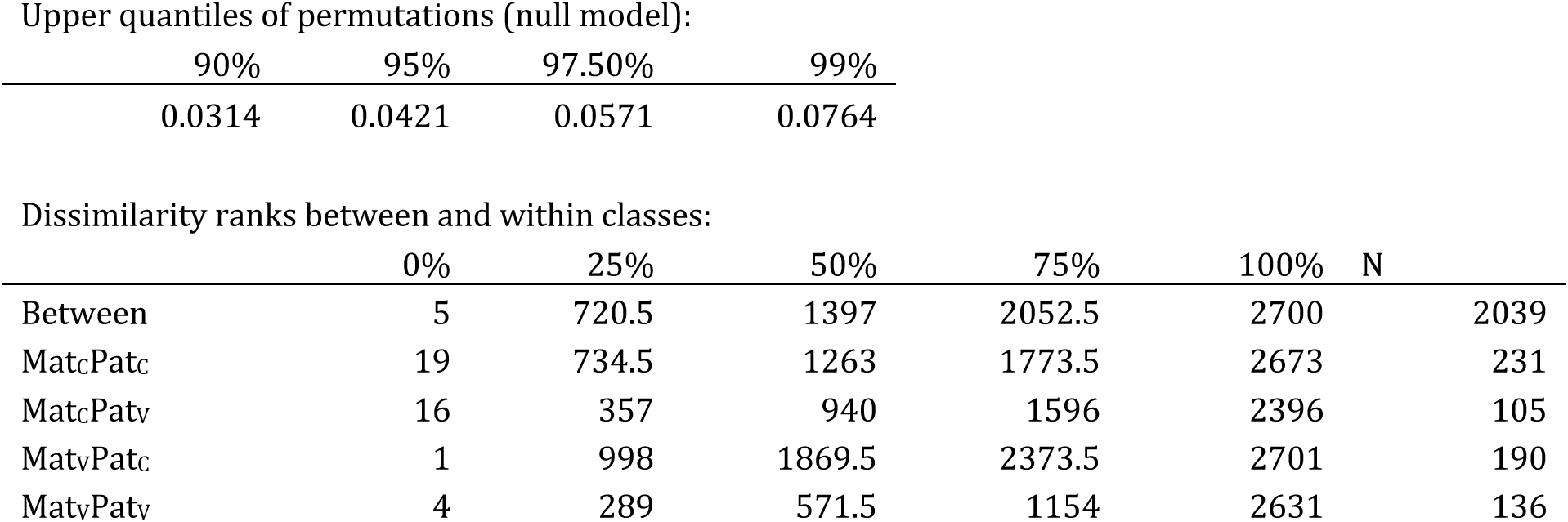
Statistical summary of ANOSIM models using a Bray-Curtis distance matrices on microbiota family counts considering only offspring samples that were kept at 18°C and either kept naïve or exposed to the experimental *Vibrio*.

| 90% | 95% | 97.50% | 99% |
| --- | --- | --- | --- |
| 0.0273 | 0.038 | 0.0508 | 0.0669 |

Dissimilarity ranks between and within classes:
|  | 0% | 25% | 50% | 75% | 100% | N |
| --- | --- | --- | --- | --- | --- | --- |
| Between | 1 | 883.5 | 1737 | 2623 | 3486 | 2635 |
| Mat<sub>c</sub>Pat<sub>c</sub> | 3 | 673 | 1585 | 2412.5 | 3401 | 231 |
| Mat<sub>c</sub>Pat<sub>v</sub> | 47 | 1153.5 | 1809 | 2465 | 3482 | 136 |
| Mat<sub>v</sub>Pat<sub>c</sub> | 2 | 594 | 1969 | 2709 | 3426 | 253 |
| Mat<sub>v</sub>Pat<sub>v</sub> | 7 | 1088.5 | 1824 | 2708.5 | 3485 | 231 |
| 0.0314 | 0.0421 | 0.0571 | 0.0764 |

Dissimilarity ranks between and within classes:
|  | 0% | 25% | 50% | 75% | 100% | N |
| --- | --- | --- | --- | --- | --- | --- |
| Between | 5 | 720.5 | 1397 | 2052.5 | 2700 | 2039 |
| MatcPatc | 19 | 734.5 | 1263 | 1773.5 | 2673 | 231 |
| MatcPatv | 16 | 357 | 940 | 1596 | 2396 | 105 |
| MatvPatc | 1 | 998 | 1869.5 | 2373.5 | 2701 | 190 |
| MatvPatv | 4 | 289 | 571.5 | 1154 | 2631 | 136 |

## Supporting Information

Supporting Data 1: Supporting Data 1_transcriptomics_preprocessing.xlsx

Supporting Data 2: Supporting Data 2_transcriptomics_final_analyses.xlsx

Supporting Data 3: Supporting Data 3_PCA_statistics.rtf

Supporting Data 4: Supporting Data 4_GO data.xlsx

Supporting Data 5: Supporting Data 5_immune cell score analysis.xlsx Supporting Protocol: Supporting_Protocol.docx

Supporting Tables: Supporting Tables.docx

## Supporting Figures

**Fig S1.**
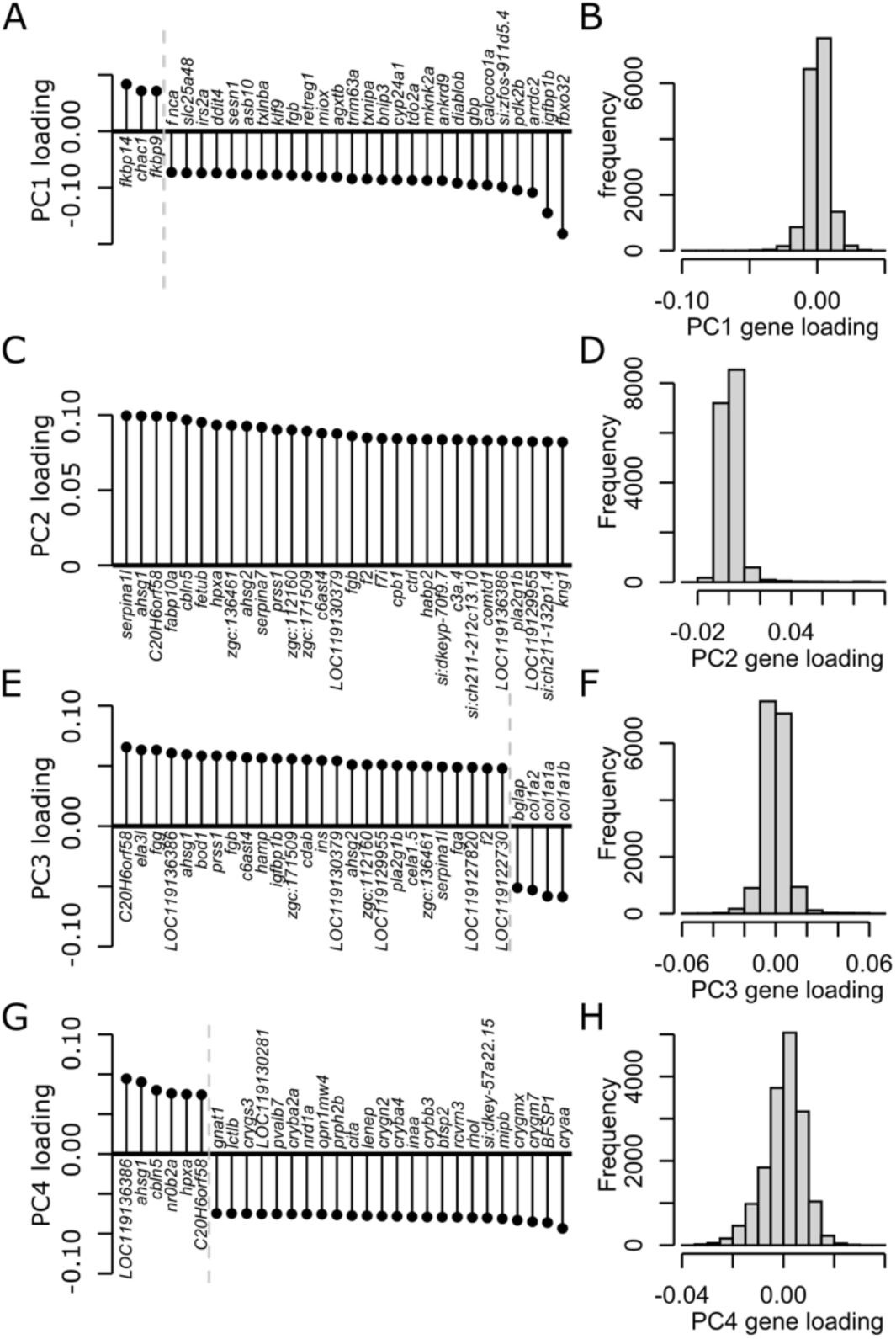
Sample scores and gene loadings from a PCA on preprocessed gene expression data. (A,C,E,G) The 30 most heavily loaded genes on PC1-PC4, respectively, and (B,D,F,H) gene loading distributions for the same PCs.

**Fig S2.**
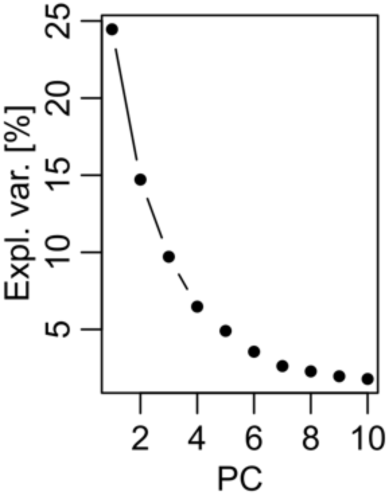
Scree-plot for the PCA on gene residuals.

**Fig S3.**
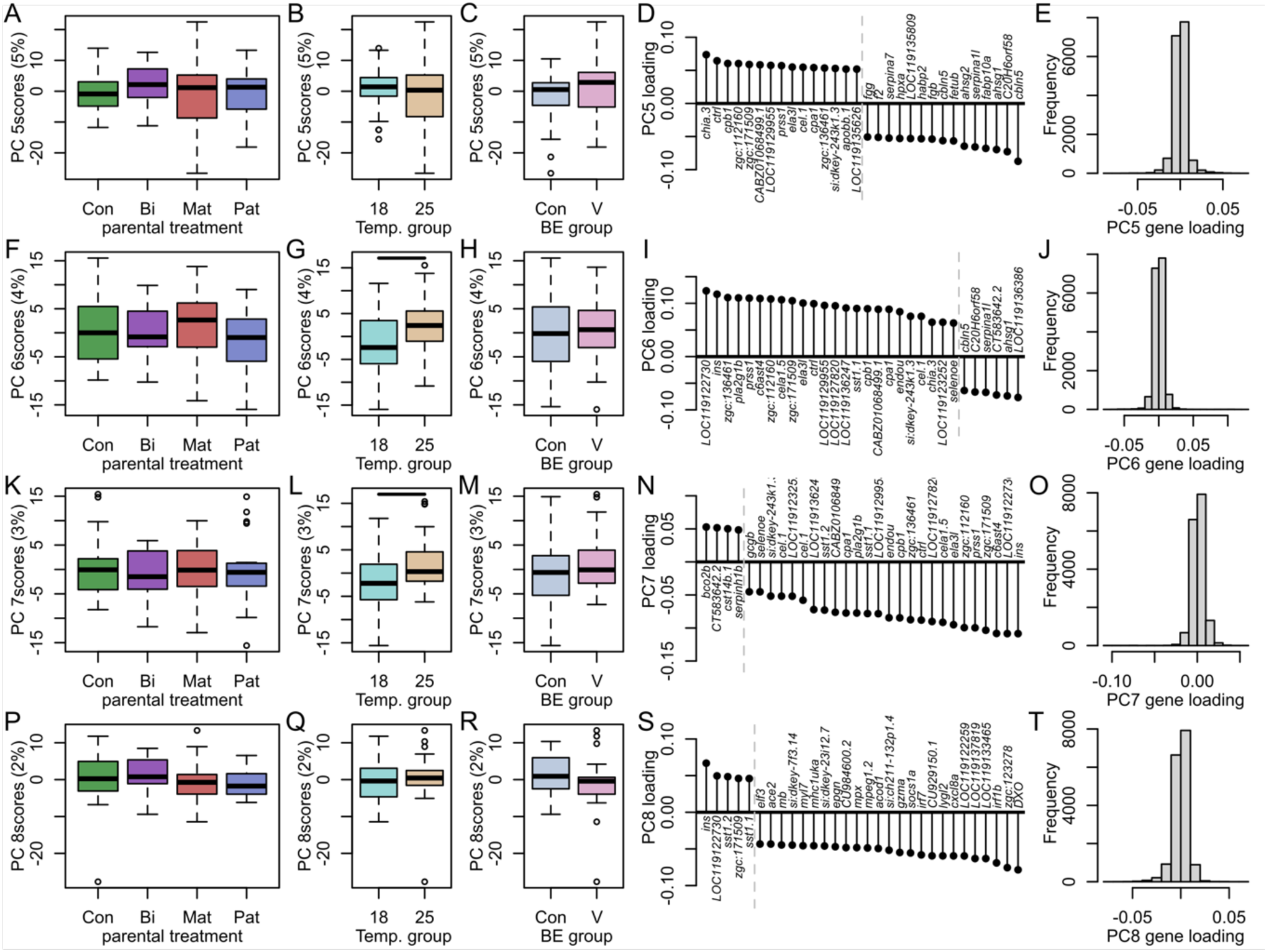
Sample scores and gene loadings from a PCA on preprocessed gene expression data. (A,F,K,P) Sample scores of PC5-PC8, respectively, for parental treatment groups. (B,G,L,Q) Samples scores for Temp treatment groups. (D,I,N,S) The 30 most heavily loaded genes on PC5-PC8, respectively, and (E,J,O,T) gene loading distributions for the same PCs. Bars indicate significant differences.

**Fig S4.**
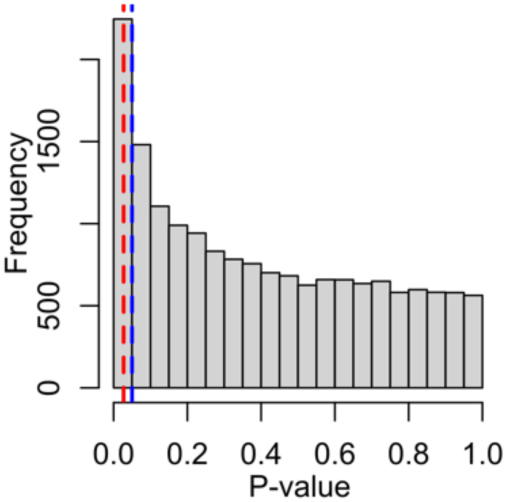
P-value histogram of *Vibrio* count and gene expression correlations. Histogram of raw p-values of the Spearman correlations between experimental *Vibrio* count and all genes’ expression profiles. The blue dashed line indicates the p=0.05% cut-off. The red dashed line indicates the p-value below which genes are deemed “significant” (p.cor=”sig”) using a custom correction method.

**Fig S5.**
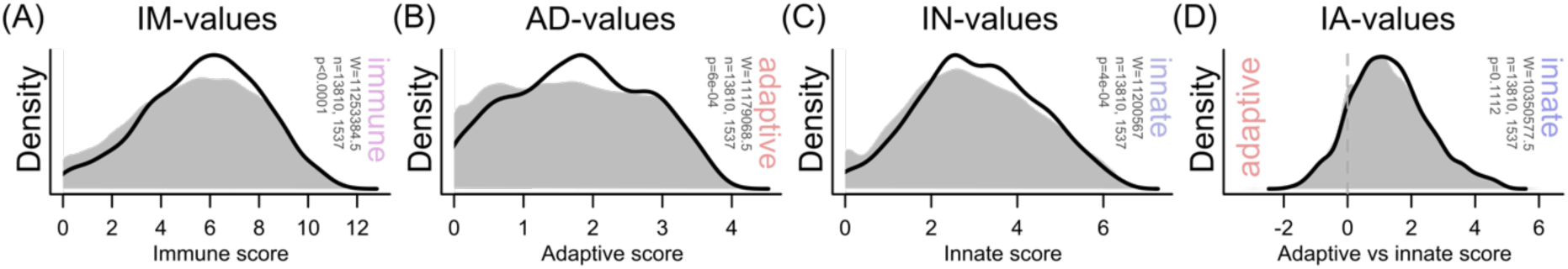
Average immune cell expression rank analysis of genes correlated to experimental *Vibrio* abundance. Distribution of IM-values (A), AD-values (B), IN-values (C) and IA-values (D) of significant genes. Significant differences in mean values between sets of significant genes and the background (all genes) were determined using Wilcoxon tests.

**Fig S6.**
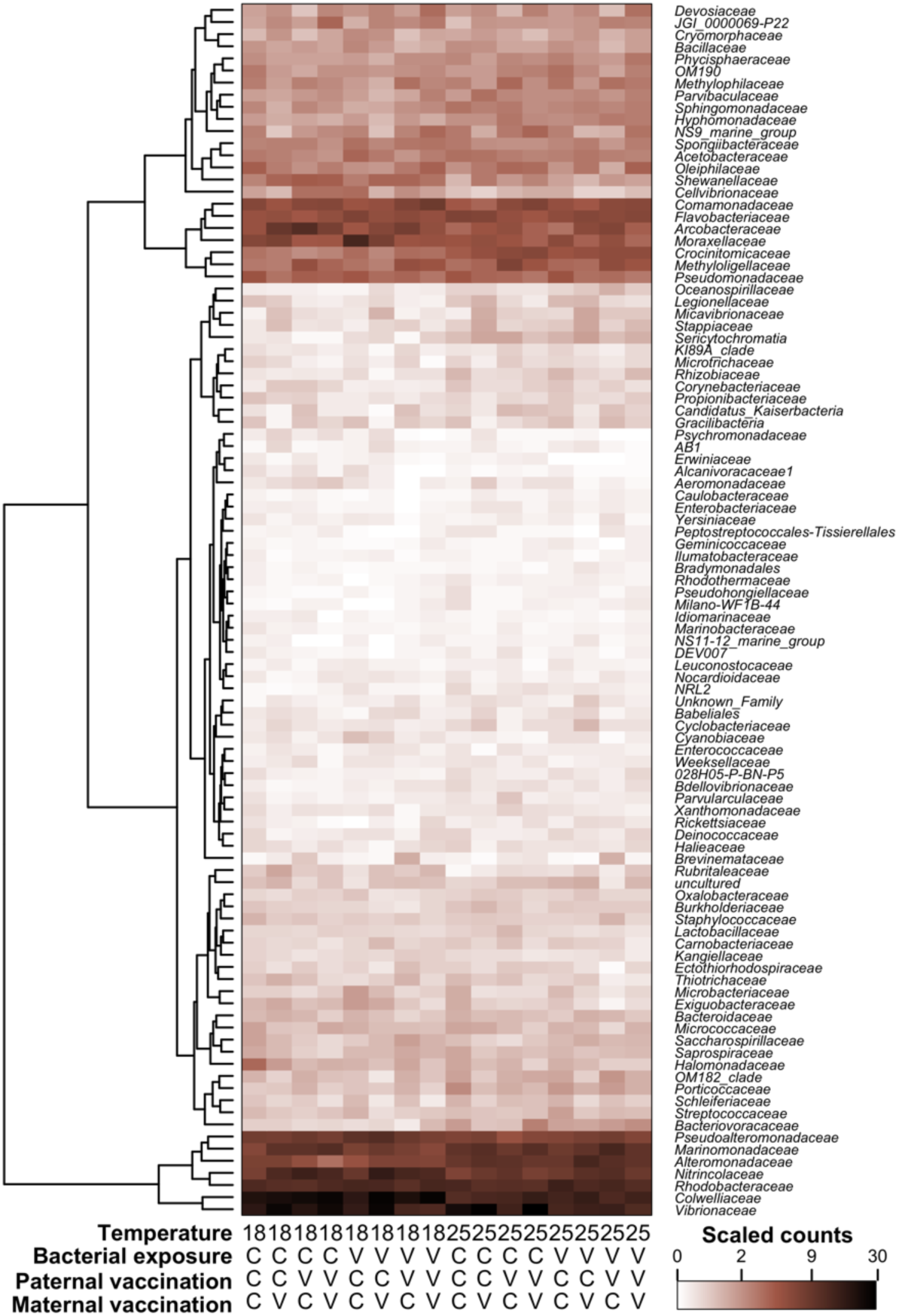
Heatmap of the top 100 microbe families with the highest scaled count sums.

**Fig S7.**
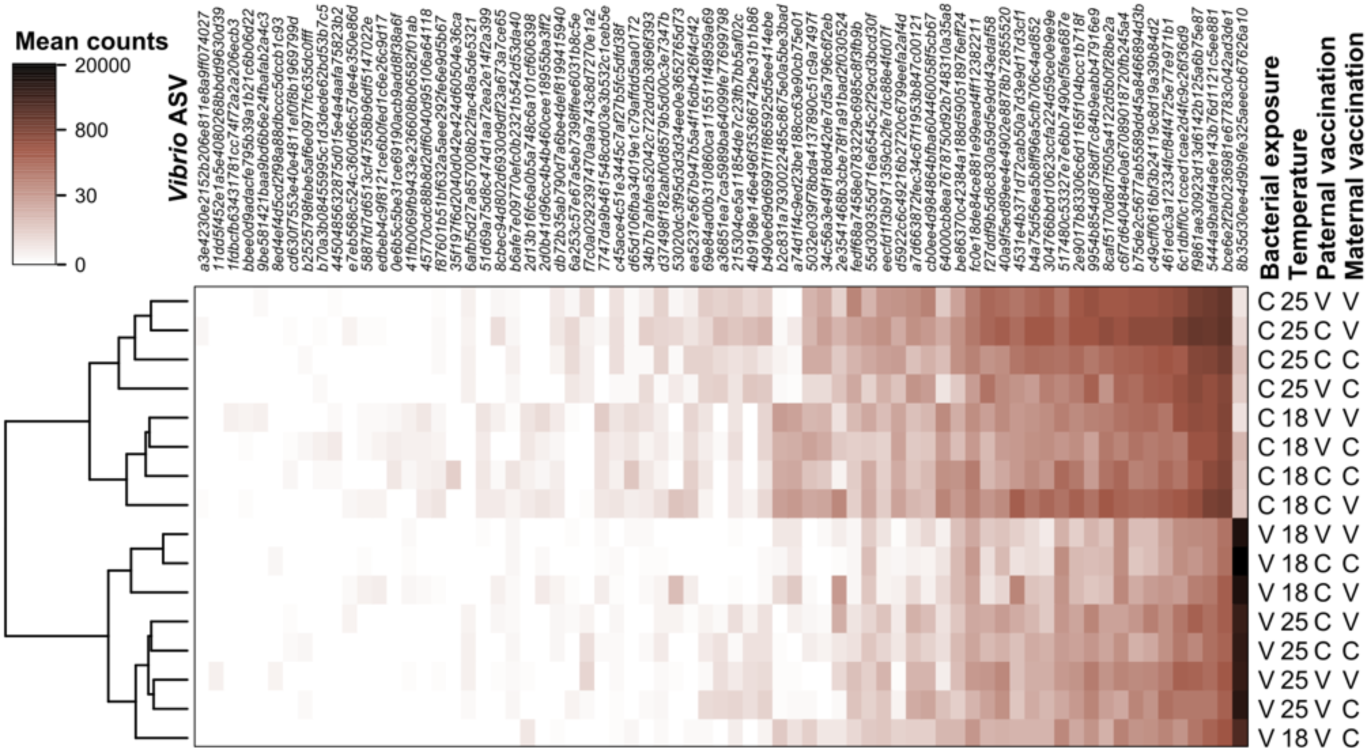
Heatmap of *Vibrio* strain counts. Strains have been sorted according to their average count inclining from left to right and only those with on average >1count/sample are shown. The strain most to the right has the highest average count and also a marked difference between samples treated with the experimental *Vibrio* strain and the control group, indicating that it is the experimental *Vibrio* strain.

**Fig S8.**
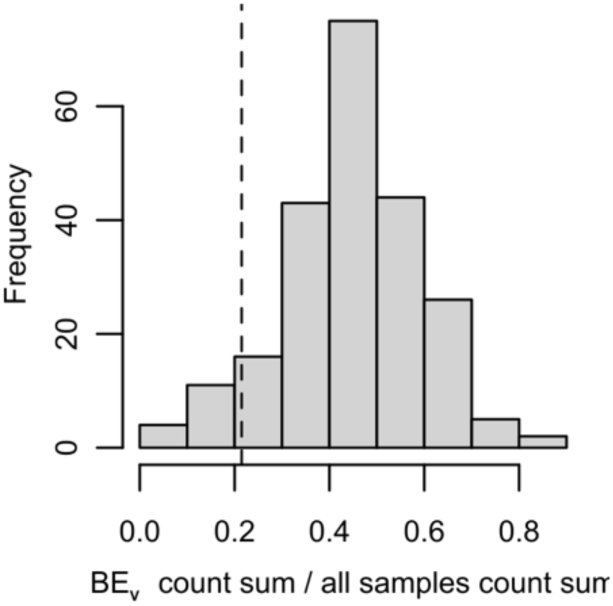
Histogram of proportions of count sums found in bacterial exposure samples per genus. The proportions across genera are normally distributed and the dashed line indicated the proportion found in the genus *Vibrio* after subtracting the experimental *Vibrio* strain.

**Fig S9.**
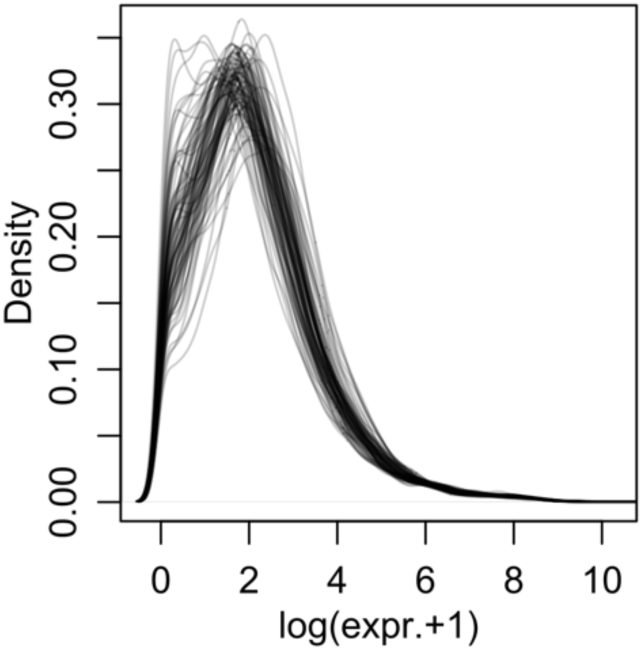
Density plots of all samples’ raw TPM values.

**Fig S10.**
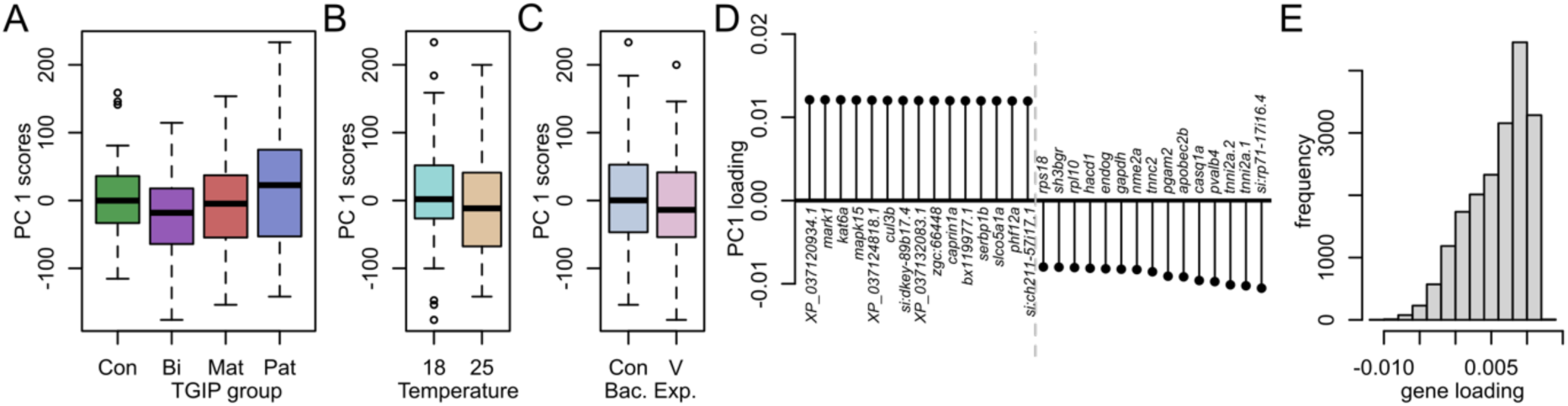
PCA plots of uncorrected TPM data. (A-C) PC1 scores did not show differentiation for any intended treatment. (D) Gene loadings on PC1 suggested that genes involved in muscle development and metabolism are highly loaded on PC1. (E) Histogram of gene loadings on PC1 showed a strongly skewed distribution.

**Fig S11.**
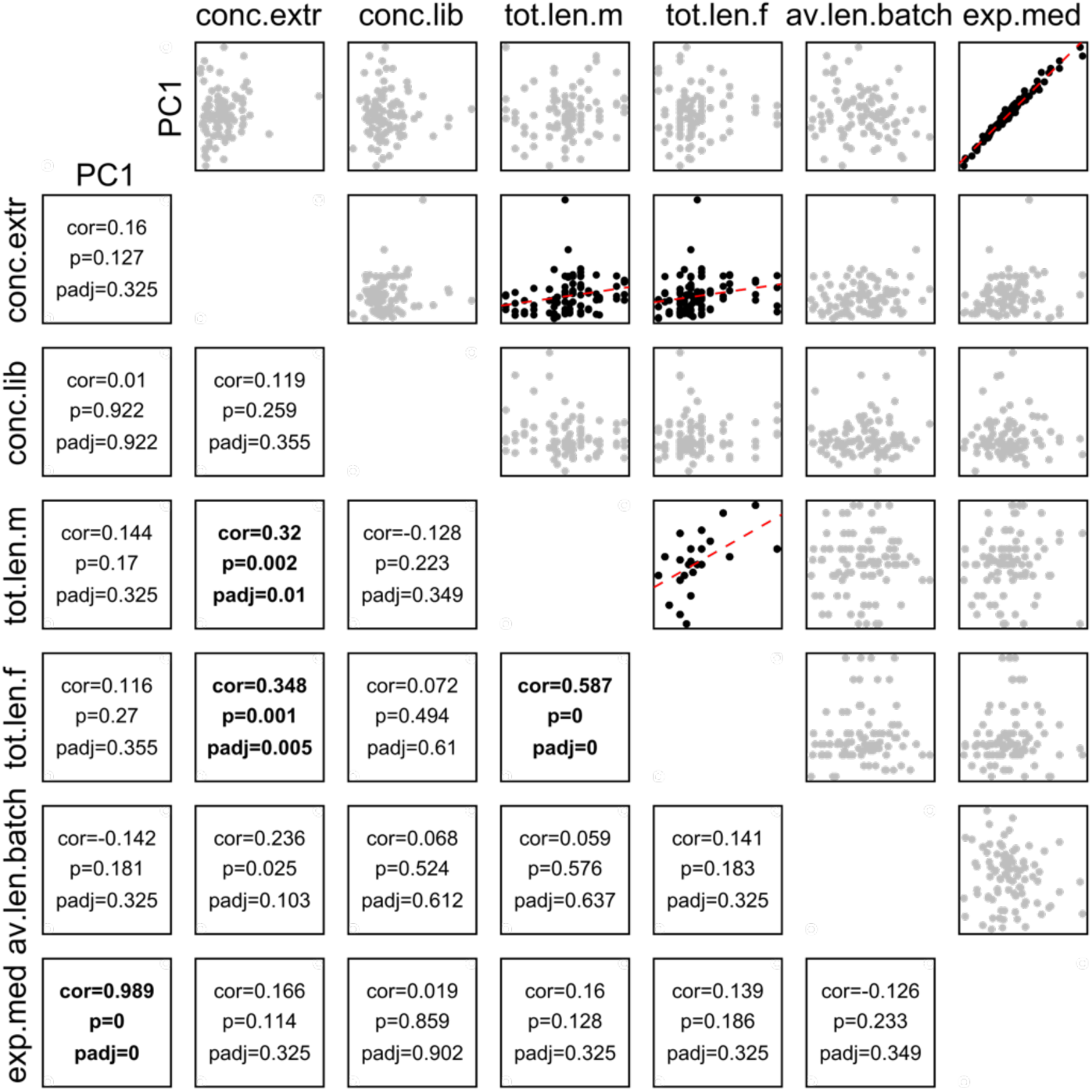
Correlations between the uncorrected PC1 and possible confounding variables. Scatterplots with regression lines and Spearman correlation test statistics for considered confounding variables with each other and the uncorrected PC1. Considered variables were: RNA concentration after extraction, RNA concentration as measured directly before library synthesis, father body length, mother body length, average body length of offspring batch, and median of samples’ expression distribution (Fig S9).

**Fig S12.**
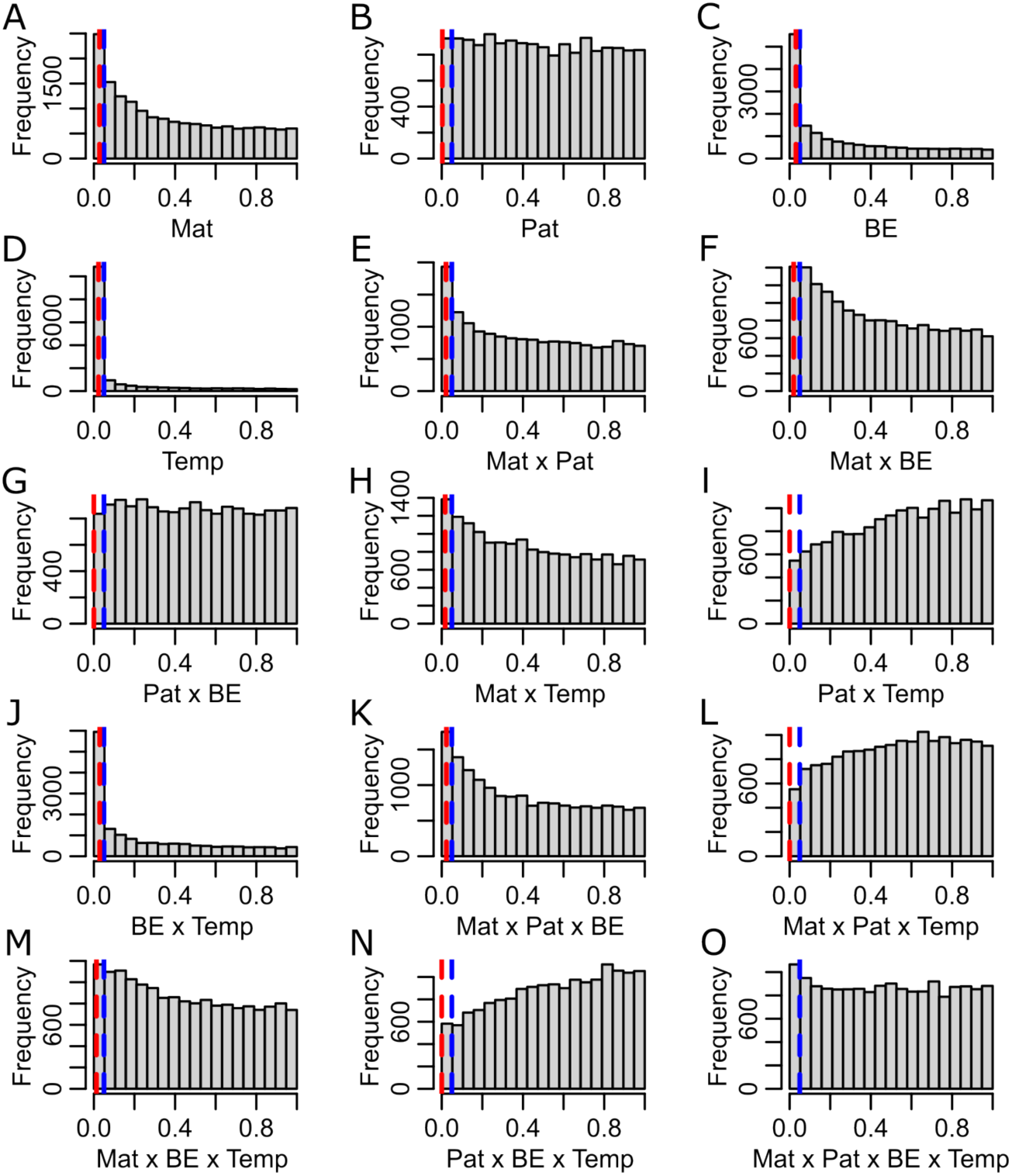
Histograms of raw p-values of the gene expression mixed model estimates. The blue dashed line indicates the 0.05% cut-off of our significance level alpha. The red dashed line indicates the p-value below which genes could be deemed “significant” (p.cor=”sig”) using a custom correction method.

