## Supporting Data 3 for "Parent-specific transgenerational immune priming enhances offspring defense – unless heat-stress negates it all"

**PC1**

**Parental treatment:**

Dunnett's test for comparing several treatments with a control :

95% family-wise confidence level

$Con

diff	lwr.ci	upr.ci	pval

Bi-Con 5.226382 -8.442220 18.89499 0.6778

Mat-Con 7.908775 -5.308593 21.12614 0.3435

Pat-Con 1.786685 -12.489705 16.06307 0.9809

---

Signif. codes: 0 '***' 0.001 '**' 0.01 '*' 0.05 '.' 0.1 ' ' 1

**Temp treatment:**

Welch Two Sample t-test

data: pcar_juv$x[which(fac == levels(fac)[1]), pc] and pcar_juv$x[which(fac == levels(fac)[2]), pc]

t = 8.5811, df = 85.011, p-value = 3.802e-13

alternative hypothesis: true difference in means is not equal to 0

95 percent confidence interval:

19.37297 31.05801

sample estimates:

mean of x mean of y

12.60775 -12.60775

**IC treatment**

Welch Two Sample t-test

data: pcar_juv$x[which(fac == levels(fac)[1]), pc] and pcar_juv$x[which(fac == levels(fac)[2]), pc]

t = 0.57111, df = 87.809, p-value = 0.5694

alternative hypothesis: true difference in means is not equal to 0

95 percent confidence interval:

-5.601347 10.118974

sample estimates:

mean of x mean of y

1.129407 -1.129407

**PC2**

**Parental treatment:**

Dunnett's test for comparing several treatments with a control :

95% family-wise confidence level

$Con

diff	lwr.ci	upr.ci	pval

Bi-Con 9.440031 -1.159563 20.03962 0.0913 .

Mat-Con 7.114098 -3.135577 17.36377 0.2334

Pat-Con 5.561336 -5.509579 16.63225 0.4814

---

Signif. codes: 0 '***' 0.001 '**' 0.01 '*' 0.05 '.' 0.1 ' ' 1

**Temp treatment:**

Welch Two Sample t-test

data: pcar_juv$x[which(fac == levels(fac)[1]), pc] and pcar_juv$x[which(fac == levels(fac)[2]), pc]

t = -3.0092, df = 85.985, p-value = 0.003436

alternative hypothesis: true difference in means is not equal to 0

95 percent confidence interval:

-14.825853 -3.029966

sample estimates:

mean of x mean of y

-4.463955 4.463955

**IC treatment**

Welch Two Sample t-test

data: pcar_juv$x[which(fac == levels(fac)[1]), pc] and pcar_juv$x[which(fac == levels(fac)[2]), pc]

t = -1.5857, df = 84.519, p-value = 0.1165

alternative hypothesis: true difference in means is not equal to 0

95 percent confidence interval:

-10.972391	1.236393

sample estimates:

mean of x mean of y

-2.434	2.434

**PC3**

**Parental treatment:**

Dunnett's test for comparing several treatments with a control :

95% family-wise confidence level

$Con

diff	lwr.ci	upr.ci	pval

Bi-Con -1.515597 -10.35539 7.324192 0.9538

Mat-Con -1.972788 -10.52075 6.575178 0.8985

Pat-Con -2.962050 -12.19491 6.270809 0.7769

---

Signif. codes: 0 '***' 0.001 '**' 0.01 '*' 0.05 '.' 0.1 ' ' 1

**Temp treatment:**

Welch Two Sample t-test

data: pcar_juv$x[which(fac == levels(fac)[1]), pc] and pcar_juv$x[which(fac == levels(fac)[2]), pc]

t = 4.5542, df = 86.574, p-value = 1.71e-05

alternative hypothesis: true difference in means is not equal to 0

95 percent confidence interval:

5.868262 14.958393

sample estimates:

mean of x mean of y

5.206664 -5.206664

**IC treatment**

Welch Two Sample t-test

data: pcar_juv$x[which(fac == levels(fac)[1]), pc] and pcar_juv$x[which(fac == levels(fac)[2]), pc]

t = -2.7279, df = 87.837, p-value = 0.007697

alternative hypothesis: true difference in means is not equal to 0

95 percent confidence interval:

-11.493788 -1.805134

sample estimates:

mean of x mean of y

-3.32473	3.32473

**PC4**

**Parental treatment:**

Dunnett's test for comparing several treatments with a control :

95% family-wise confidence level

$Con

diff	lwr.ci	upr.ci	pval

Bi-Con 1.811114 -5.230834 8.853063 0.8669

Mat-Con 3.760861 -3.048616 10.570337 0.4066

Pat-Con 5.467639 -1.887436 12.822714 0.1878

---

Signif. codes: 0 '***' 0.001 '**' 0.01 '*' 0.05 '.' 0.1 ' ' 1

**Temp treatment:**

Welch Two Sample t-test

data: pcar_juv$x[which(fac == levels(fac)[1]), pc] and pcar_juv$x[which(fac == levels(fac)[2]), pc]

t = -0.33302, df = 87.827, p-value = 0.7399

alternative hypothesis: true difference in means is not equal to 0

95 percent confidence interval:

-4.765149 3.397342

sample estimates:

mean of x mean of y

-0.3419518 0.3419518

**IC treatment**

Welch Two Sample t-test

data: pcar_juv$x[which(fac == levels(fac)[1]), pc] and pcar_juv$x[which(fac == levels(fac)[2]), pc]

t = -3.5063, df = 87.471, p-value = 0.0007194

alternative hypothesis: true difference in means is not equal to 0

95 percent confidence interval:

-10.588917 -2.927469

sample estimates:

mean of x mean of y

-3.379096 3.379096

**PC5**

**Parental treatment:**

Dunnett's test for comparing several treatments with a control :

95% family-wise confidence level

$Con

diff	lwr.ci	upr.ci	pval

Bi-Con	2.2503233 -4.036428 8.537074 0.7179

Mat-Con -0.4062340 -6.485444 5.672976 0.9969

Pat-Con -0.7904956 -7.356793 5.775802 0.9829

---

Signif. codes: 0 '***' 0.001 '**' 0.01 '*' 0.05 '.' 0.1 ' ' 1

**Temp treatment:**

Welch Two Sample t-test

data: pcar_juv$x[which(fac == levels(fac)[1]), pc] and pcar_juv$x[which(fac == levels(fac)[2]), pc]

t = 1.0313, df = 80.333, p-value = 0.3055

alternative hypothesis: true difference in means is not equal to 0

95 percent confidence interval:

-1.729858 5.451580

sample estimates:

mean of x mean of y

0.9304305 -0.9304305

**IC treatment**

Welch Two Sample t-test

data: pcar_juv$x[which(fac == levels(fac)[1]), pc] and pcar_juv$x[which(fac == levels(fac)[2]), pc]

t = -1.6225, df = 88.813, p-value = 0.1082

alternative hypothesis: true difference in means is not equal to 0

95 percent confidence interval:

-6.4577516 0.6522318

sample estimates:

mean of x mean of y

-1.45138	1.45138

**PC6**

**Parental treatment:**

Dunnett's test for comparing several treatments with a control :

95% family-wise confidence level

$Con

diff	lwr.ci	upr.ci	pval

Bi-Con -0.5133224 -5.908221 4.881576 0.9913

Mat-Con 0.6469484 -4.569851 5.863748 0.9814

Pat-Con -2.4067887 -8.041576 3.227999 0.6033

---

Signif. codes: 0 '***' 0.001 '**' 0.01 '*' 0.05 '.' 0.1 ' ' 1

**Temp treatment:**

Welch Two Sample t-test

data: pcar_juv$x[which(fac == levels(fac)[1]), pc] and pcar_juv$x[which(fac == levels(fac)[2]), pc]

t = -2.6002, df = 89, p-value = 0.01091

alternative hypothesis: true difference in means is not equal to 0

95 percent confidence interval:

-6.9032002 -0.9228942

sample estimates:

mean of x mean of y

-1.956524 1.956524

**IC treatment**

Welch Two Sample t-test

data: pcar_juv$x[which(fac == levels(fac)[1]), pc] and pcar_juv$x[which(fac == levels(fac)[2]), pc]

t = -0.58075, df = 87.47, p-value = 0.5629

alternative hypothesis: true difference in means is not equal to 0

95 percent confidence interval:

-3.999893 2.190888

sample estimates:

mean of x mean of y

-0.4522513 0.4522513

**PC7**

**Parental treatment:**

Dunnett's test for comparing several treatments with a control :

95% family-wise confidence level

$Con

diff	lwr.ci	upr.ci	pval

Bi-Con -1.3690897 -6.011339 3.273160 0.8155

Mat-Con -0.1699413 -4.658939 4.319056 0.9994

Pat-Con -0.8167546 -5.665426 4.031917 0.9560

---

Signif. codes: 0 '***' 0.001 '**' 0.01 '*' 0.05 '.' 0.1 ' ' 1

**Temp treatment:**

Welch Two Sample t-test

data: pcar_juv$x[which(fac == levels(fac)[1]), pc] and pcar_juv$x[which(fac == levels(fac)[2]), pc]

t = -2.995, df = 89.333, p-value = 0.003551

alternative hypothesis: true difference in means is not equal to 0

95 percent confidence interval:

-6.329682 -1.280856

sample estimates:

mean of x mean of y

-1.902634 1.902634

**IC treatment**

Welch Two Sample t-test

data: pcar_juv$x[which(fac == levels(fac)[1]), pc] and pcar_juv$x[which(fac == levels(fac)[2]), pc]

t = -1.7003, df = 88.048, p-value = 0.09261

alternative hypothesis: true difference in means is not equal to 0

95 percent confidence interval:

-4.8360879 0.3763665

sample estimates:

mean of x mean of y

-1.11493	1.11493

**PC8**

**Parental treatment:**

Dunnett's test for comparing several treatments with a control :

95% family-wise confidence level

$Con

diff	lwr.ci	upr.ci	pval

Bi-Con	1.5235783 -2.792671 5.839828 0.7260

Mat-Con -0.6199984 -4.793758 3.553761 0.9690

Pat-Con -0.9492049 -5.457380 3.558971 0.9199

---

Signif. codes: 0 '***' 0.001 '**' 0.01 '*' 0.05 '.' 0.1 ' ' 1

**Temp treatment:**

Welch Two Sample t-test

data: pcar_juv$x[which(fac == levels(fac)[1]), pc] and pcar_juv$x[which(fac == levels(fac)[2]), pc]

t = -0.38318, df = 89.371, p-value = 0.7025

alternative hypothesis: true difference in means is not equal to 0

95 percent confidence interval:

-2.961500 2.003898

sample estimates:

mean of x mean of y

-0.2394003 0.2394003

**IC treatment**

Welch Two Sample t-test

data: pcar_juv$x[which(fac == levels(fac)[1]), pc] and pcar_juv$x[which(fac == levels(fac)[2]), pc]

t = 1.9779, df = 84.81, p-value = 0.05119

alternative hypothesis: true difference in means is not equal to 0

95 percent confidence interval:

-0.01281132 4.85562403

sample estimates:

mean of x mean of y

1.210703 -1.210703
