## Supporting Protocol for "Parent-specific transgenerational immune priming enhances offspring defense – unless heat-stress negates it all"

***(Pre-)analysis of the gene expression data set***

A TPM (transcripts per million) expression table of 92 juvenile samples was imported into R (v. 4.1.3,(1)) for downstream analyses and genes that were expressed in fewer than three samples were discarded, leaving 19,055 genes (Supporting Data 1: sheet “raw tpm”). Density plots of gene expression profiles were generated for each individual (Fig. S9), revealing rather heterogeneous expression patterns, which may be indicative of non-homologous tissue sampling.

To explore main axes of variation in the gene expression data set, a Principal Component Analysis (PCA) was used on the log(gene_expr+1) transformed, centered and scaled gene expression data set (Fig. S10, Supporting Data 1: sheets “original PCA…”). When investigating the scores and loadings of PC1, we noticed that none of the major treatments seemed to be reflected on PC1 scores (Fig. S10A-C), using Dunnett’s Test with “Con” as reference level to test for differences among parental treatments, and t-tests to test for differences between Temp and BE levels (all p>0.1; package “DescTools “, v. 0.99.47;(2)). Additionally, genes heavily negatively loaded on this PC unexpectedly appeared to be associated with energy metabolism (and, to a lesser degree, muscle tissue; Fig. S10D), and a histogram of the loadings suggested an atypical negative skew, indicating that PC1 might reflect unintended variation present in the data set (Fig. S10E).

A set of possible technical and biological covariates were investigated for correlations with PC1, which would suggest an influence on overall gene expression patterns (Fig. S11). These included per sample (i) the total RNA concentration after extraction and (ii) after shipping (as determined by the sequencing company), (iii) the respective juvenile’s father and (iv) mother total body length, (v) the average body length of juveniles of the respective batch (i.e., “beaker”), and (vi) the median of gene expression across all genes (which reflects the variation in gene expression profiles mentioned above, Fig. S9). These variables and PC1 were correlated to each other using Spearman Rank-sum correlations and obtained p-values were corrected using the false-discovery-rate (FDR) method. The results revealed that PC1 scores were highly significantly associated with the median of the log gene expression (ρ=0.99, p.adj<0.001), indicating that the variation in gene expression profiles displayed in Fig. S10 indeed is the conceptually unintended major source of variation in the data-set. To understand the cause of this variation, PC1 loadings were inspected. The large majority of genes (16,979) showed positive loading, while much fewer (2,076) showed negative loading (Fig. S10E). The 1% of most positively loaded genes (n=191) were analyzed for Gene Ontology (GO) biological process enrichment. The human GO annotation was used as it is the most comprehensive, even though not for all genes human ortholog annotations were present (n=165 with a human gene annotation; n=157 of those with GO annotation) and their function in human (and thus GO association) may be different compared to fish (corresponding results for GO analysis using *Danio rerio* annotations is reported in Supporting Data 4, but remained less conclusive and thus is not discussed here). We tested for GO enrichment using the gost(…,ordered_query=F, organism="hsapiens", significant=T, correction_method="gSCS", sources=c("GO:BP"), custom_bg=all_expressed_genes, domain_scope="custom") function (“gprofiler2” v.0.2.1;(3); Supporting Data 1: sheet “raw tpm” & Supporting Data 4: sheets “raw PC1…”; all_expressed_genes include 14,435 human orthologs). Eight terms were found significantly enriched, of which four were cellular regulation/organization related, while the remaining were generic regulatory processes, such as chromatin organization (Supporting Data 4; sheet “PC1 pos load hsapiens”). When this was done with the 1% most negatively loaded genes (n=132 of which had a human ortholog and n=121 of these a GO annotation), twenty-one processes were identified and almost all appeared to be related to energy metabolism (Supporting Data 4; sheet “PC1 neg load hsapiens”). These GO terms, the overall skew in gene loading frequencies towards positive loadings (Fig. S10E) and the common energy metabolism genes with negative loading (Fig. S10D; also, several muscle-related GO terms narrowly missed significance) suggest that variation reflected in PC1 is linked to differences in tissue composition (possibly reflecting an varying amount of the functionally distinct foregut portion of the gut) of samples, which is also reflected in heterogeneous expression profiles (Fig. S9), possibly as a result of inconsistencies during gut dissections.

To correct for this effect and to identify whether and how the intended treatment factors affected gene expression, an unconventional analysis approach was chosen over more orthodox pipelines to remove unintended variation in gene expression reflected in this PC1 in our subsequent statistical analysis. In detail, individual genes were investigated and a gene was removed from the pipeline if it was not expressed in at least all but one individual of at least two treatment groups (from among the 16 treatment combinations), and mean log(gene expression+1) levels among individuals expressing the gene had to be at least 0.1 to be deemed meaningful. In this manner, 1,567, 7, and 832 genes were excluded for being too low expressed, being expressed in too few samples, or both, respectively, leaving 16,649 to be considered in the following steps (see Supporting Data 1 for TPM tables, model estimates, significance testing). Then, for each gene, a linear model was computed, using aforementioned log-transformed gene expression as dependent variable and PC1 scores as dependent variable and its residuals were stored (Supporting Data 1: sheet “tpm PC1 residuals”).

*Final PCA and pairwise comparisons of gene expression*

Gene residuals were used to compute two gene-wise linear mixed models (package “nlme”, v. 3.1-161; (4,5)). As multiple offspring came from distinct families, the family factor was expected to affect not only our statistical analysis, but also ordination analyses. Thus, a first linear mixed model was calculated per gene, using no independent variables and only a random intercept to obtain a second set of residuals (corrected now also for the family effect; lme(log_gene_expression_residuals ~ 1, random = ~1|family, method="REML"); Supporting Data 2: sheet “tpm family residuals for PCA”). These residuals were stored but exclusively used to compute a final PCA (Fig. 2, Fig. S1-3). A second mixed model per gene included also treatment groups and their interactions as independent factors (model in R: lme(log_gene_expression_residuals ~ Mat * Pat * BE * Temp, random = ~1|family, method="REML"); Supporting Data 1: sheets “mixed model…”). The Anova() function was used to produce type II ANOVA tables (package “car”, v. 3.0;(6); raw p-values, fdr-corrected p-values and a custom significance evaluation (see below) are reported in aforementioned Supporting Data 1 sheets).

The number of significant estimates from the linear mixed model computed for each gene is roughly in line with identifying Temp and BE treatments, as well as their interaction, as the major effects on gene expression (see Fig. 2; Table S1; fdr-corrected significant p-values: 10,243, 3,116 and 3,924 significant estimates, respectively). Other main effects or interactions produced only very few to no fdr-corrected significant estimates, but we noted that for several variables and interactions many more raw p-values below 0.05 across genes were observed than expected by chance, suggesting that fdr correction might have over-corrected these raw p-values (Table S1, Fig. S12; see also http://varianceexplained.org/statistics/interpreting-pvalue-histogram/). To circumvent this, for each estimate, we calculated the proportion of raw p-values below 0.05 and compared this to our expectation of a proportion of 0.05 (as α=0.05), assuming no effect. While we did find that for some independent variables indeed the proportion of genes with a raw p-value below 0.05 was approximately 0.05 (those with flat p-value histogram column distributions; see Fig. S12), for others a substantially higher proportion of genes was found to have p-values below 0.05, despite none of them attained significance after fdr correction (those with higher columns in the corresponding p-value range in the p-value histogram column distributions; see Fig. S12). For instance, we found 2,389 genes with a raw p-value <0.05 for the Mat factor although only 833 (corresponding to 5% of tested genes) were expected under H_0_ assumptions - but still only 2 retained significance after fdr correction (Fig. S12: red dashed line corresponds to 833 genes with a raw p<0.05). Thus, we expected to find about 1,556 genes to be considered “significant” after *post hoc* correction, as these exceed the expected number of 833 genes under the assumption that there was no effect (Table S1). Due to this discrepancy in p-value number, and the suggested parental effects revealed by the PCA (especially in interactions; Fig. 2), we used pairwise comparisons between focal treatment groups to evaluate treatment effects (Fig. 3). Still, per independent variable we also report a number of the most significant genes corresponding to this “expected” number of *post hoc* corrected significant genes (i.e., number of raw significant genes minus 833; “p_cor_=yes”; Table 1; Supporting Data 1: sheet “Mixed model 18°C”) and provide GO enriched terms (if any; Supporting Data 4: sheets “est…”).

For pairwise comparisons the function glht() (package “multcomp”, v.1.4-19;(7)) was used to test seven treatment groups (Mat_V_Pat_C_BE_C_Temp_18_, Mat_C_Pat_V_BE_C_Temp_18_, Mat_C_Pat_C_BE_V_Temp_18_, Mat_C_Pat_C_BE_C_Temp_25_, Mat_V_Pat_C_BE_V_Temp_18_, Mat_C_Pat_V_BE_V_Temp_18_ and Mat_V_Pat_V_BE_V_Temp_18_) against a reference group (Mat_C_Pat_C_BE_C_Temp_18_) based on the mixed model, and estimates and raw p-values were computed for each pairwise comparison (Supporting Data 2: “pairwise 18°C” sheets). Also, to compute corresponding comparisons with the reference temperature of 25°C, the aforementioned procedure was repeated with the reference level of the Temp factor being set to 25°C (Supporting Data 2: “pairwise 25°C” sheets). Raw p-values were corrected for multiple testing using the p.adjust() function and the “fdr” method per comparison across all genes (Supporting Data 2: sheets “pairwise …°C p-fdr-values”).

*Average immune cell expression rank analyses*

To explore if sets of significantly expressed genes are enriched for genes with pronounced expression in immune cells, a reference single cell expression data set based on human data was downloaded from the human protein atlas website using the R package “HPAanalyze” (v. 1.16.0;(8)). The “RNA single cell type” data set was chosen, which contains normalized expression data for 79 different annotated cell types (incl. “undifferentiated”) collated from various studies’ data sets. Of these cells, 13 were identified as being immune cell related, and while “dendritic cells” were not assigned to one of the two branches of the immune system, those specific to the innate branch were "granulocytes", "Hofbauer cells", "Kupffer cells", "Langerhans cells", "macrophages", “microglial cells” and "monocytes", while those specific to the adaptive branch were "B-cells", "NK-cells" (natural killer cells), "Plasma cells", "T-cells" and “thymic epithelial cells”. All 14,435 human orthologs from our study were matched by Ensembl ID to the genes within the downloaded data base. For each of these genes, we aimed to estimate its association in expression (in humans) to immune cell types, i.e, a high association reflects that a gene is predominantly expressed across immune cell types, and rarely or not at all in other cell types. For this, first, cell types with no gene expression were excluded and the remaining cells were sorted according to their average normalized expression values (in human) and ranks were assigned (the cell type with the highest expression value obtained the highest rank). Obtained rank numbers were then divided by the number of non-zero cells and the rank sum of the designated immune cells (IM value), those assigned to the innate (IN value), and adaptive immune system (AD value) were calculated, resulting in three values that reflect how strong a given gene’s expression of is associated with cells of the respective type relative to all others (higher values indicate stronger association). Additionally, to illustrate if a gene is associated rather with cell types of one branch of the immune system over the other, the rank sums of the adaptive cell types were subtracted from the rank sum of the innate cell types, resulting in a value (IA) in which more positive numbers indicate higher gene expression association with cell types of the innate immune system, while more negative values indicate this for cell types of the adaptive immune system (Supporting Data 5). While these values by themselves are not very informative, their distribution across all genes of the background gene set served as reference to investigate the distribution of these values in sets of target genes, e.g., sets of differentially expressed genes. To determine if target gene sets contained genes that on average had different IM, IN, AD or IA values compared to the reference background, Wilcoxon signed rank tests were performed and the distribution of ranks was plotted for both backgrounds and target gene sets (Fig. 4; Fig. S5).

***Microbiome - additional analyses***

Our analyses indicated that counts belonging to *Vibrio* strains among the sequenced 16s rRNAs are in sum more abundant in BE_V_ samples compared to BE_C_ samples. Considering the high abundance of the experimental *Vibrio* strain relative to all other *Vibrio* strains (Fig. S6-7), we tested whether this effect may be an artifact caused by limited sequencing depth (i.e., if a large proportion of read depth was claimed by experimental *Vibrio*, relatively fewer reads might have represented remaining *Vibrio* and *non-Vibrio* strains). However, an alternative explanation may be that the experimental *Vibrio* strain suppressed/outcompeted other (*Vibrio*) strains with similar ecology specifically due to their large number. To gauge the potential influence of these two effects, first, per sample we calculated the total number of *Vibrio* counts across all strains and subtracted the counts of the experimental strain. Then, we divided the sum of counts from samples that were exposed the experimental bacteria (BEV) by the sum of those which were not (BE_C_), which provided a gauges the count bias mentioned before. We identified that only approximately 21.5% of non-experimental *Vibrio* counts were found in samples that were exposed to experimental bacteria. To understand if this an atypical percentage across microbiota genera, for each sample we calculated the sum of all counts of strains belonging to each genus identified in this study (853 genera). Then, for each genus, we calculated the proportion of counts found in samples exposed to the experimental *Vibrio*. We excluded genera that had counts in less than 5% of the 307 individuals considered, and we excluded the genus *Vibrio*, leading to 226 genera considered. We determined that aforementioned ratio was significantly lower in *Vibrio* than the average among other genera (one-sample Wilcoxen Signed-Rank Test: V = 25132, n = 226, p-value < 0.0001; after normality of the ratio distribution among other genera could not be confirmed: Shapiro-Wilk: W = 0.98643, p-value = 0.03, Fig. S8), suggesting that the effect of non-experimental *Vibrio* strains being underrepresented in the bacterial exposure individuals is likely, at least partially, due to the experimental strain suppressing or outcompeting other *Vibrio* strains, which may occupy similar niches.
